# EEG Source Identification through Phase Space Reconstruction and Complex Networks

**DOI:** 10.1101/2020.09.08.287755

**Authors:** Morteza Zangeneh Soroush

## Abstract

Artifact elimination has become an inseparable part while processing electroencephalogram (EEG) in most brain computer interface (BCI) applications. Scientists have tried to introduce effective and efficient methods which can remove artifacts and also reserve desire information pertaining to brain activity. Blind source separation (BSS) methods have been receiving a great deal of attention in recent decades since they are considered routine and standard signal processing tools and are commonly used to eliminate artifacts and noise. Most studies, mainly EEG-related ones, apply BSS methods in preprocessing sections to achieve better results. On the other hand, BSS methods should be followed by a classifier in order to identify artifactual sources and remove them in next steps. Therefore, artifact identification is always a challenging problem while employing BSS methods. Additionally, removing all detected artifactual components leads to loss of information since some desire information related to neural activity leaks to these sources. So, an approach should be employed to suppress the artifacts and reserve neural activity. In this study, a new hybrid method is proposed to automatically separate and identify electroencephalogram (EEG) sources with the aim of classifying and removing artifacts. Automated source identification is still a challenge. Researchers have always made efforts to propose precise, fast and automated source verification methods. Reliable source identification has always been of great importance. This paper addresses blind source separation based on second order blind identification (SOBI) as it is reportedly one of the most effective methods in EEG source separation problems. Then a new method for source verification is introduced which takes advantage of components phase spaces and their dynamics. A new state space called angle space (AS) is introduced and features are extracted based on the angle plot (AP) and Poincare planes. Identified artifactual sources are eliminated using stationary wavelet transform (SWT). Simulated, semi-simulated and real EEG signals are employed to evaluate the proposed method. Different simulations are performed and performance indices are reported. Results show that the proposed method outperforms most recent studies in this subject.

## 1. Introduction

EEGs, which contain brain electrical activity, are recorded through a non-invasive technique by means of electrodes located on the scalp [1]. EEGs have become very effective signals in terms of diagnosis mental disorders like epilepsy, autism, depression and etc. These nonlinear and non-stationary signals can be employed to study cognitive states of human mind. Thanks to use of EEG in brain computer interfaces (BCI), disabled individuals are more likely to control different machines and devices such as computers, wheelchairs and etc. [9]. Now it is possible to study and analyze sleep disorders through EEG processing [21]. Unfortunately, in most practical settings EEGs are usually corrupted by environmental and physiological signals called EEG artifacts. Biological artifacts, including electromyogram (EMG), electrocardiogram (ECG), electrooclugram (EOG), eye blinking artifact and etc., levitate from non-cerebral sources in human body. In contrast, environmental noises and artifacts arise from external sources such as line power transmission, electric motors, electrode movement and etc. [13]. Both types of artifacts interfere with EEG signals easily and make interpretation and diagnosis difficult. Non-physiological artifacts are precluded by most EEG recording devices but biological artifacts like EMG and EOG still remain and need to be eliminated. This fact motivates us to study and propose a new method in this paper to reduce biological artifacts. As it was mentioned above, EEGs have a wide usage in different fields so interpreting corrupted EEGs is of a great deal of importance. Accurate signal classification and precise recognition are totally dependent on artifacts and noises. Needless to say, artifact removal and noise suppression are inseparable parts in biological signal processing and the more effective the methods are the more precise and accurate the results will be. Therefore numerous studies have been conducted to eliminate noise and artifacts from EEGs [13–16]. There are several methods to deal with corrupted EEGs such as linear filtering, autoregressive modeling, adaptive filters, blind source separation (BSS) based methods, wavelet transforms, principal component analysis (PCA) and etc. [1–8]. Conventional methods like linear filters are not appropriate and effective due to inherent overlap in frequency domain between artifacts and cerebral activity. BSS based methods have been receiving a great deal of attention in EEG processing and artifact suppression since they isolate noise and artifacts into independent components (ICs) using subspace filtering [13]. SOBI algorithm utilizes the original EEG and time shifted version(s) in order to exploit temporal information and estimate uncorrelated components. SOBI has been employed in several studies to remove artifacts [10–12]. Artifact removal based on BSS methods consists of three major steps: (i) applying the source separation method, (ii) source identification and artifact removal, (iii) channel reconstruction using mixing matrix and remaining sources. Experimental simulations prove that these methods are practical in many cases. Based on previous studies, these methods are useful tools in EEG processing [2–4]. Several studies have been carried out in order to investigate these methods with the aim of removing artifacts [10–12, 15–17]. Different articles have come to a conclusion that Independent Component Analysis (ICA), introduced as a noise suppression tool for the first time in [18], is the most robust method in artifact elimination but is not very time fast. Among different BSS based methods, second order blind identification (SOBI) is reportedly the most effective one. SOBI has been employed to remove artifacts in several studies. Several authors have found SOBI the most reliable and widely used approach [2–7, 22, 23]. Several toolboxes like EEGLAB [20] have implemented SOBI due to its wide usage and efficiency. SOBI has been known as a superior method in comparison with ICA and most BSS methods. Therefore we decide to use SOBI in this study to extract EEG sources. More detail information about SOBI is brought in following sections. To achieve reliable results, extracted sources should be identified in order to eliminate artifacts and noise. Sources used to be visually identified by experts. In this method, artifactual sources are classified and removed by experts but most of the time this method leads to insufficient EEG data for further analysis. Moreover, in some cases the origin of the artifacts or noises are not known. Thus source identification should be applied to achieve reliable neural sources. Manual identification methods are time consuming and expensive. Additionally, in real time and practical projects, manual methods seem to be useless. Researchers have tried to propose automated methods to identify extracted sources [10, 11 and 12]. Mostly, sources are identified by classifiers based on their features and characteristics [9, 13]. Several features have been proposed by previous studies. Since biological signals are nonlinear, chaotic features and nonlinear analysis seem to be more likely to be successful in artifact removal and noise reduction [14]. This motivates us to examine phase space of the extracted sources in order to verify them. Phase space reconstruction is usually performed to transfer a signal or time series into a new space in order to analyze the behavior of the given signal. Most nonlinear and chaotic features like correlation dimension (CD), fractal dimension (FD), Lyaponouv exponents (LE), recurrence quantification analysis (RQA) and etc. are extracted through the phase space **[REF]**. In 1901, W. Gibbs introduced and explained phase space reconstruction for the first time **[REF]**. This analysis is able to demonstrate signal characteristics. This study aims to analyze characteristic of the sources via the phase space. It is believed that artifacts and noise affect the amplitude of the signal considerably. So we tend to ignore source amplitude and consider phase information. With this regard, we introduce a new state space extracted from the phase space of the given signal. This new space is based on the angle values between points in the phase space and called angle space which results in a graphical illustration named angle plot. Moreover, Poincare planes are claimed to be very effective ways to describe nonlinear signals based on their representation in a state space, angle plot for example. Since AP is a graphical representation of the underlying signal, Poincare planes are employed to describe and quantify the AP. Suggested features are extracted based on AP and Poincare planes. Extracted features are normalized and then sources are classified using common and conventional classifiers such as multilayer perceptron (MLP) neural network, K nearest neighbor (KNN) and Bayes. We also apply the mixture of these classifiers to improve the results. Identified sources as artifacts, are fed into the artifact removal procedure which is based on SWT. Several studies have claimed the advantages to usage of SWT due to its abilities in terms of non-stationary and nonlinear signals [54]. We employ SWT to prevent data loss since there is always information leakage to artifact components while using BSS methods. SWT can keep cerebral activity and reject artifacts to high extent. Remained components are used to reconstruct clean EEG. We are of the opinion that the proposed method is capable of identifying neural sources and artifacts. Not only is this method able to verify sources precisely but also it can suppress artifacts effectively. In this study, we take advantage of SOBI and AP of the extracted sources to separate and then identify them automatically. Fig 1 shows the block diagram of the suggested method. Contaminated EEGs are separated into sources via SOBI algorithm. Estimated sources are reconstructed in phase space. We propose a method to extract dynamic of the phase space for each source and then quantify it through Poincare sections. In other words, reconstructed phase space is transferred into a new space called Angle Space (AS) and some quantifiers such as Poincare intersections are defined to describe phase space dynamics mathematically. Extracted features are fed into basic classifiers to identify sources. Real and simulated signals are used in this study to assess performance of the suggested method. Different criteria like classification performance (CP), relative root-mean-square error (RRMSE), Correlation Analysis (CA) and average mutual information (AMI) are defined to evaluate this method. Results show that the proposed method is successful and effective. Graphical illustrations are presents in order to clarify the upsides of the suggested method. Typical and quantitative results are reported and explained. Also processing time is considerably low. More detail information is provided in following sections.

**Fig 1.**
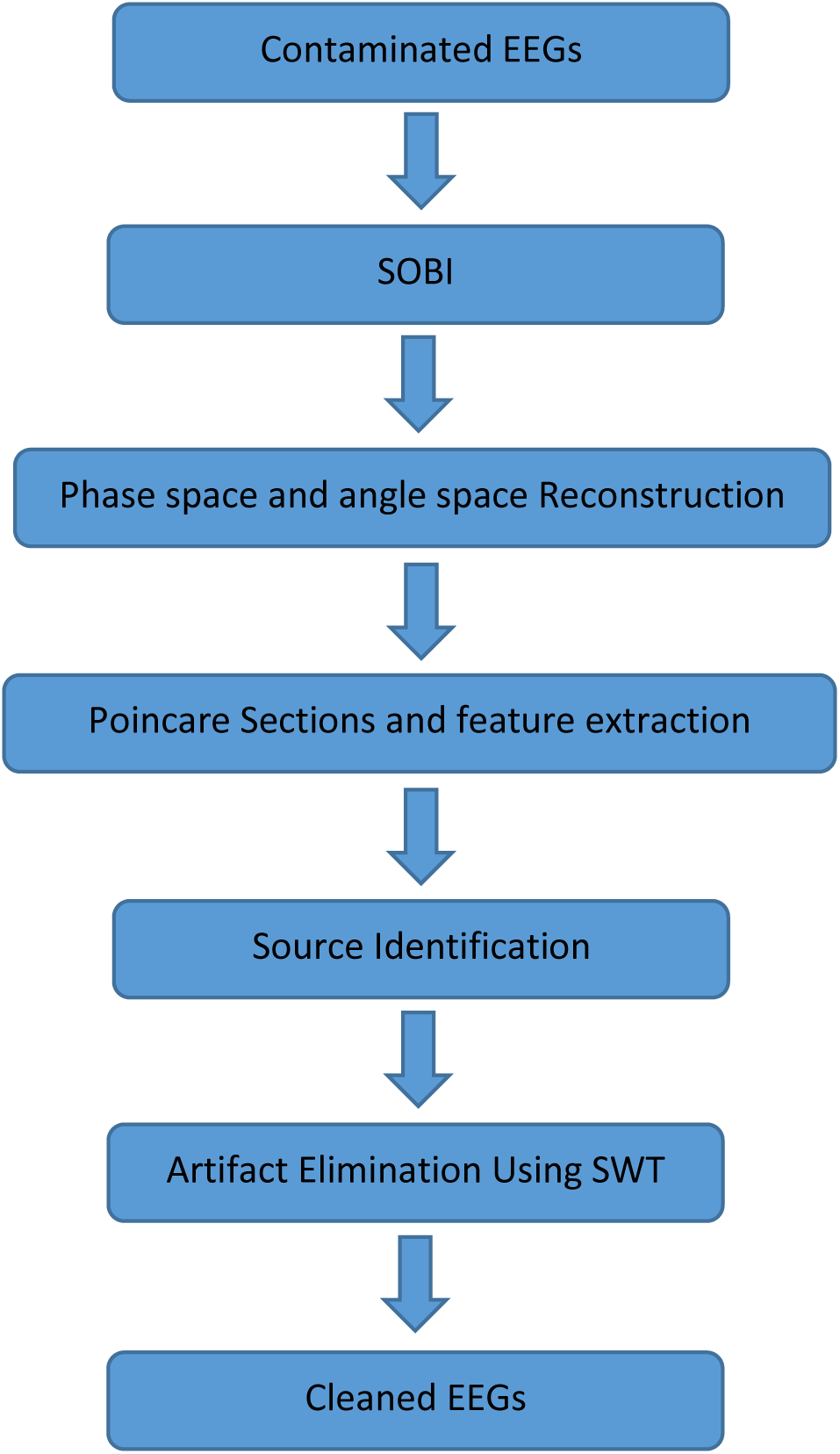
The block diagram of the proposed method

This paper is organized as follows: “Section 2” represents material and methods. In “Section 3” you can find results. “Section 4” is dedicated to discussion and finally the paper is concluded in “Section 5”.

## 2. Materials and Methods

### 2.1. Blind Source Separation and SOBI

The BSS makes effort to solve Eq. (1).

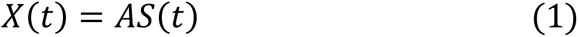

Where *X*(*t*) = {*x*_1_(*t*), …, *x*_*N*_(*t*)} and *S*(*t*) = {*s*_1_(*t*), …, *s*_*M*_(*t*)} represent observation signals for *N* channels (e.g. EEGs) and *M* estimated sources respectively. *A* is called the mixing matrix and has the size of *N* ∗ *M*. In this model, EEGs are considered instantaneous linear mixture of sources through an unknown mixing matrix of *A* [28].

SOBI algorithm is based on second order statistics and consists of two main stages: (i) signals (i.e., EEGs) are zero-meaned and whitening process is performed and (ii) a set of covariance matrices is constructed. Belouchrani et al. in [19] proposed SOBI for extracting correlated sources based on joint approximate diagonalization of a random set of time-lagged covariance matrices. Covariance matrix is defined based on Eq. (2).

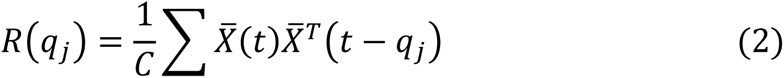

Where 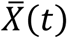 and 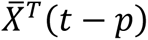 are zero-meaned and time-delayed signals respectively and *q* indicates time lags which are chosen a set of different values instead of a single time lag to improve time-efficiency of SOBI. *C* is the number of considered time lags. Sources are supposed mutually uncorrelated and stationary. In some studies it is reported that SOBI is capable of functionally separating sources which are physiologically interpretable [24–27]. SOBI is really robust in low SNRs [27–32]. Since SOBI is iterative, it is found one of the fastest algorithms in comparison with other BSS methods [28]. Comparing to ICA, SOBI relies on second order statistical analysis of signals while ICA is based on higher order statistics, which means ICA is more time consuming, complex and laborious [32]. These features suggest that SOBI method of source separation is effective and feasible. These characteristics motivated us to use SOBI in this study. For further information about SOBI algorithm and its utility refer to [29–31].

### 2.2. Phase space reconstruction

Phase space reconstruction (PSR), part of chaos theory, has become a very useful and powerful tool in nonlinear signal processing. Phase space reconstruction has been part of numerous studies **[REF]**. This powerful analysis introduces a new transformation by retaining magnitude and phase information of signals. Several characteristics of a given signal can be described through the concept of phase space. This motivated us to study these characteristics with the goal of automated source identification. In 1901, phase space was introduced by W. Gibbs for the first time **[REF]**. Phase space includes state vectors describing the signal. There are several ways to reconstruct the phase space of a signal. Having reviewed previous studies, we turn to the most common method which is time delay embedding [39, 40]. Suppose that *v*(*t*) is a signal with *K* time samples. We can reconstruct *K* − *d* + 1 vectors in the phase space as:

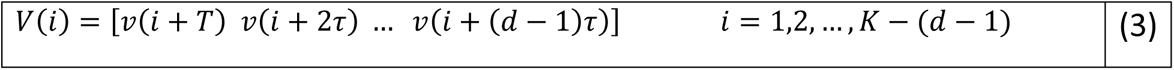

Where *d* and *τ* are the embedding dimension and time delay respectively. *d* and *τ* are important parameters while reconstructing phase space. Several studies have been conducted to estimate these parameters [37, 38]. Based on previous studies, which are related to EEG processing through phase space, the value of *d* is chosen as two and *τ* is 0.2-times the standard deviation of the signal [34–36].

### 2.3. Angle space reconstruction

Having reconstructed phase space of the signal, we consider the angle between each three points (in row) as a geometrical characteristic of the phase space. In other words, each line connecting points in the phase space is considered a vector. The angles between vectors and also the vector length are calculated in order to transform the phase space into a new state space called angle space (AS). Fig 2 shows the process of reconstructing angle space from a two dimensional phase space. Angle space reconstruction leads to two sequences of angle values (AV) and vector lengths (VL) which contain valuable information about the underlying signal.

**Fig 2.**
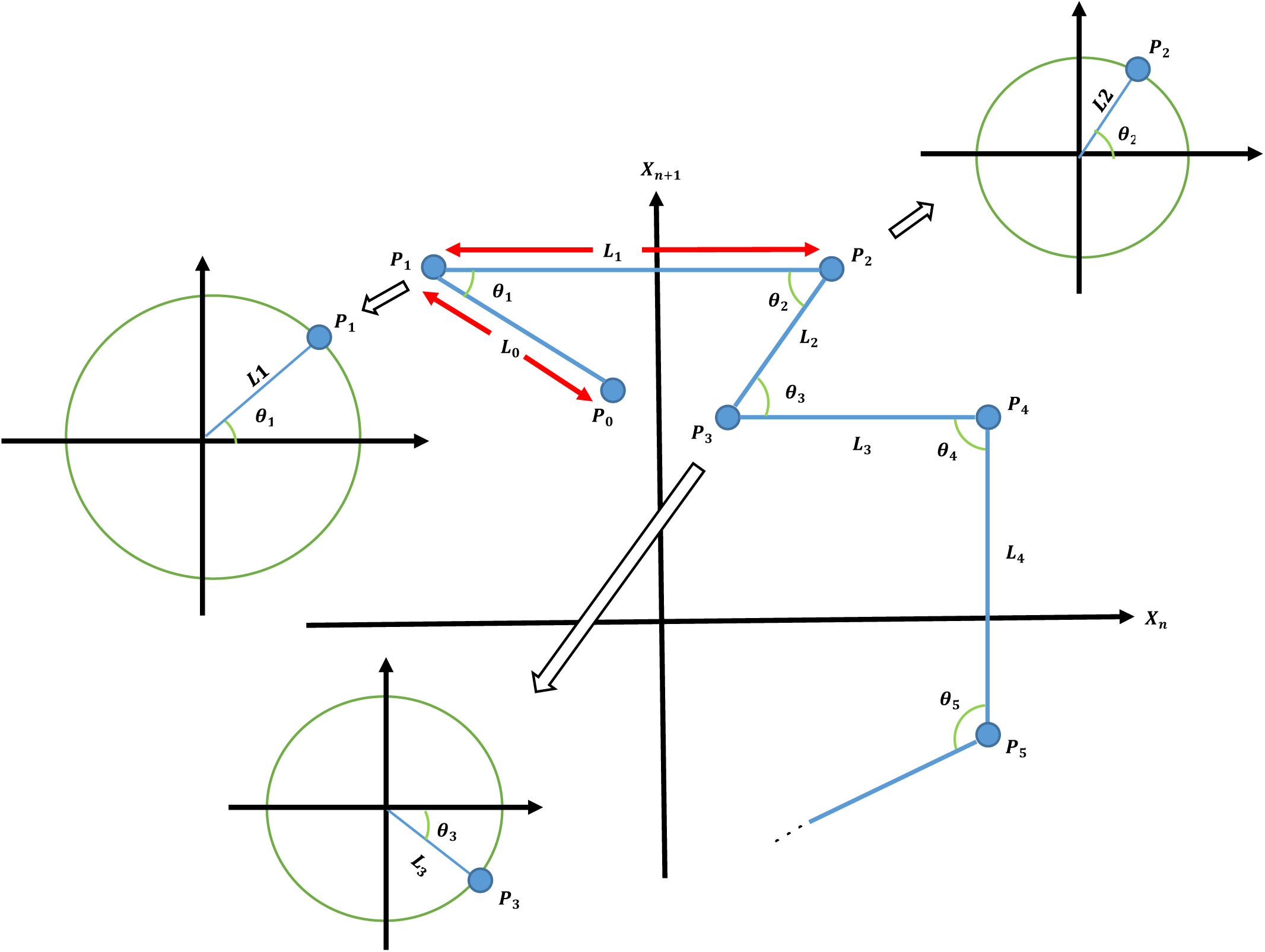
Phase space of a hypothetical signal. Angle and vector length are considered for all points in the phase space except for the first and last ones.

In Fig 2, *P*_0_, *P*_1_, *P*_2_, …, *P*_*K*−*d*_ are the points in the phase space of a hypothetical signal. The first and last points are ignored. *θ*_1_, *θ*_2_, *θ*_3_, …, *θ*_*K*−*d*−1_ and *L*_1_, *L*_2_, *L*_3_, …, *L*_*K*−*d*−1_ are two sequences which are estimated based on the phase space and are necessary to reconstruct the angle space. The main goal of this subsection is to introduce the angle space and its characteristics. By this new description, signals seem possible to be classified according to their representation in angle space. Fig 3 illustrates AS and estimated angle and length sequences for the hypothetical phase space given in Fig 2.

**Fig 3.**
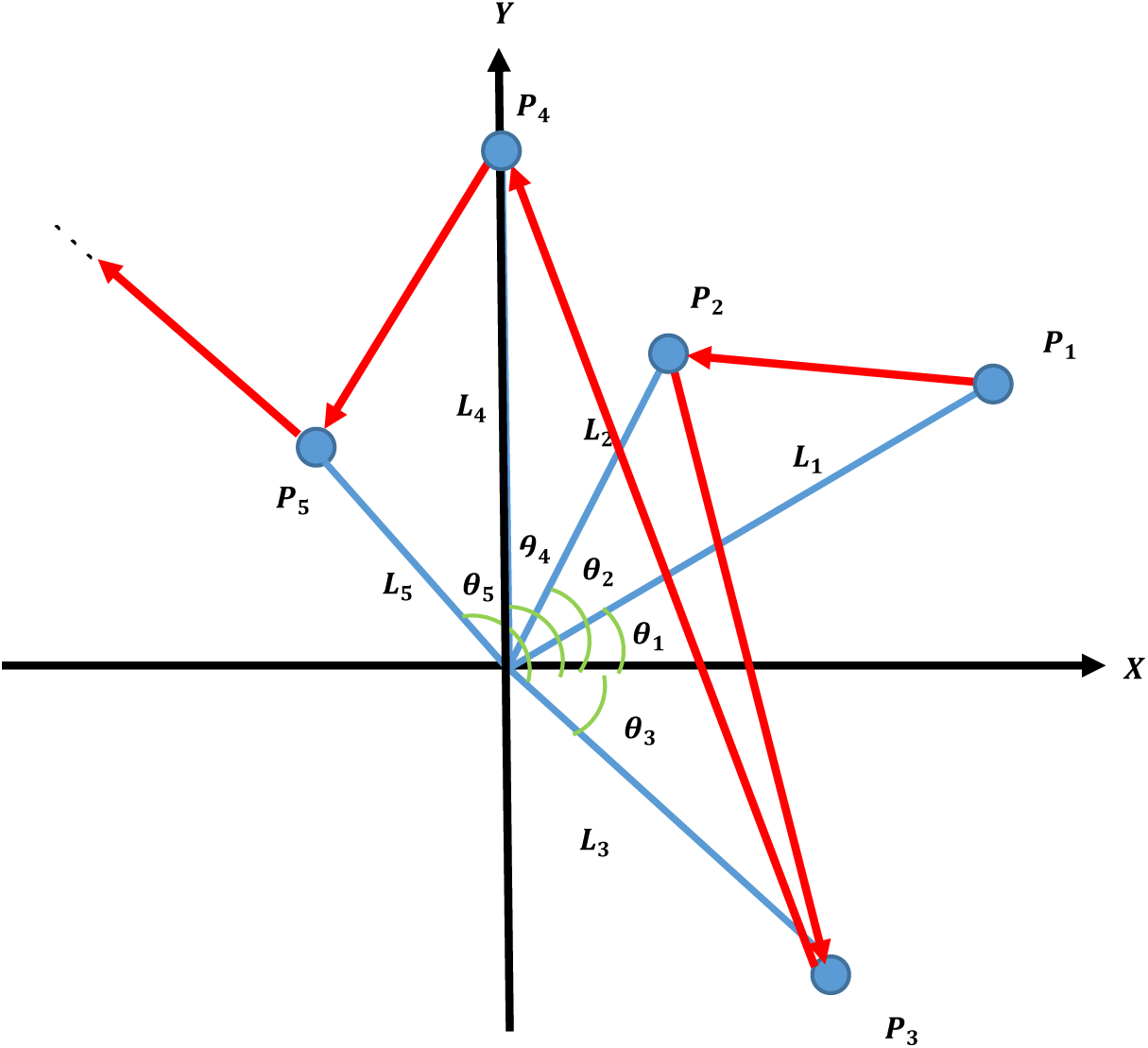
AS for the given signal in Fig.2.

Since signal amplitude is mostly affected by artifacts and noise, we aim to consider signal phase as the only source of information in this study. We believe that signal phase is rich enough to be used in further steps. So signal length is set to unit for all points in AS in order to achieve AP. Therefore we suppose the vector length equal to one and all angle values are transferred to the X-Y coordination on the unit circle in order to study angle space and its dynamics. We here decide to ignore vector length which is correspondent to signal amplitude. Fig 4 demonstrates the illustration of angle values on the unit circle (*r* = 1). We call this representation angle plot (AP). It can be considered a new representation of a signal.

**Fig 4.**
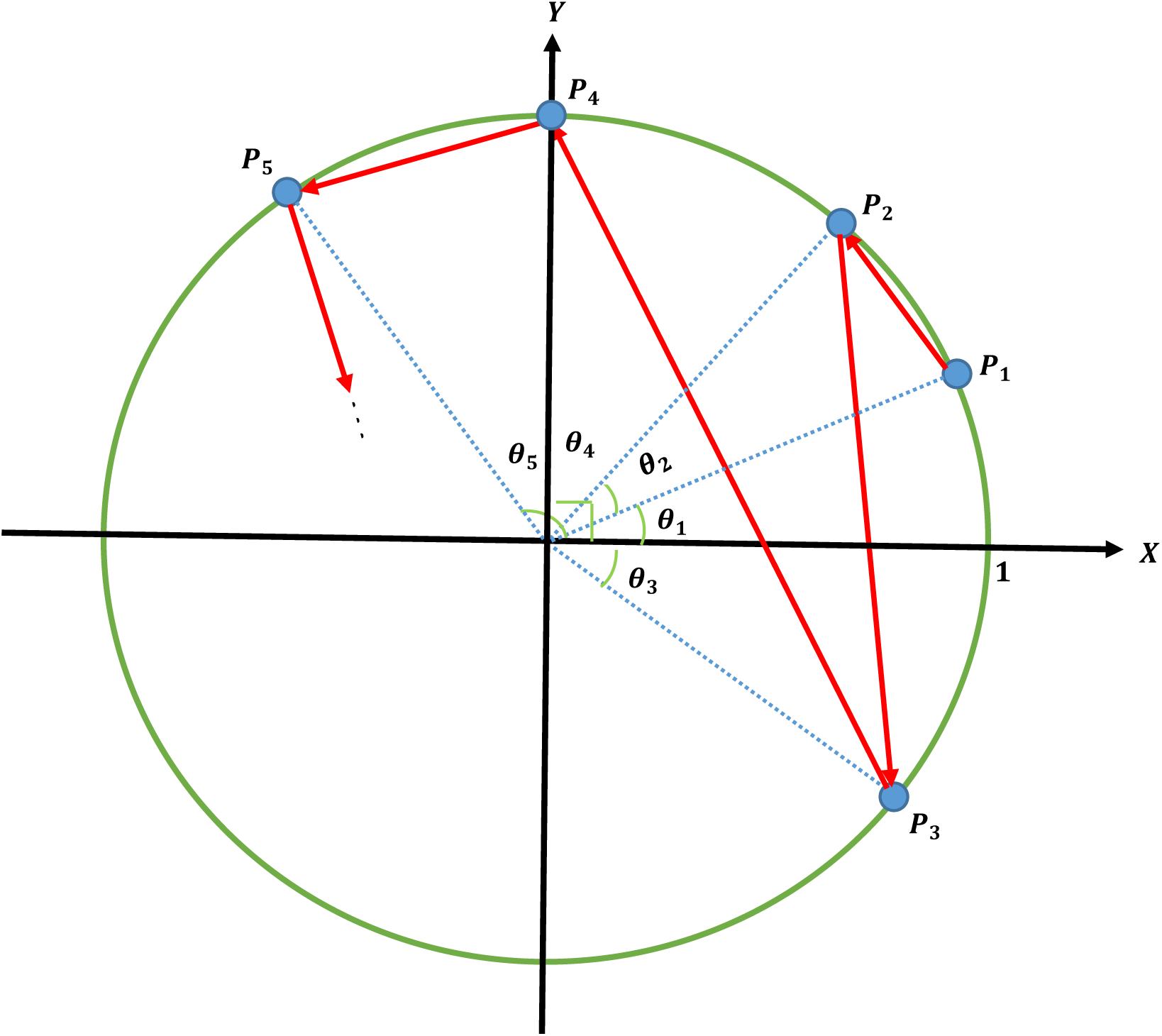
AP for the given phase space in Fig 2

Nonlinear and dynamical systems contain two main parts which are state and dynamic. Both parts should be analyzed to achieve essential information of the system and its changes through the time [39]. In this study, angle values with connecting lines are plotted to represent both state and dynamics of the extracted sources. This analysis provides us with a new graphical representation for signals which is used in following steps. Different features are defined and then extracted from this new representation.

### 2.4. Feature extraction based on AP and Poincare Planes (PPs)

#### 2.4.1. Poincare plane

Poincare sections are considered as geometrical description of a state space. PPs are defined in one dimensional less than the corresponding state space. These planes were first introduced by Poincare and then studied by many researchers. PPs enable us to analyze signal trajectories and transitions. In other words, PPs are effective tools to study system dynamics. Therefore we made a decision to apply this technique. Choosing appropriate PPs is of great deal of importance. Thanks to suitable PPs, maximum information about system dynamics and changes is transferred and also down sampled [45]. Having reviewed previous studies, we came to the conclusion to employ five suggested PPs [46]. We call these five sections PP1 to PP5. Fig 5 shows the AP for a hypothetical signal explained above and the Poincare sections.

**Fig 5.**
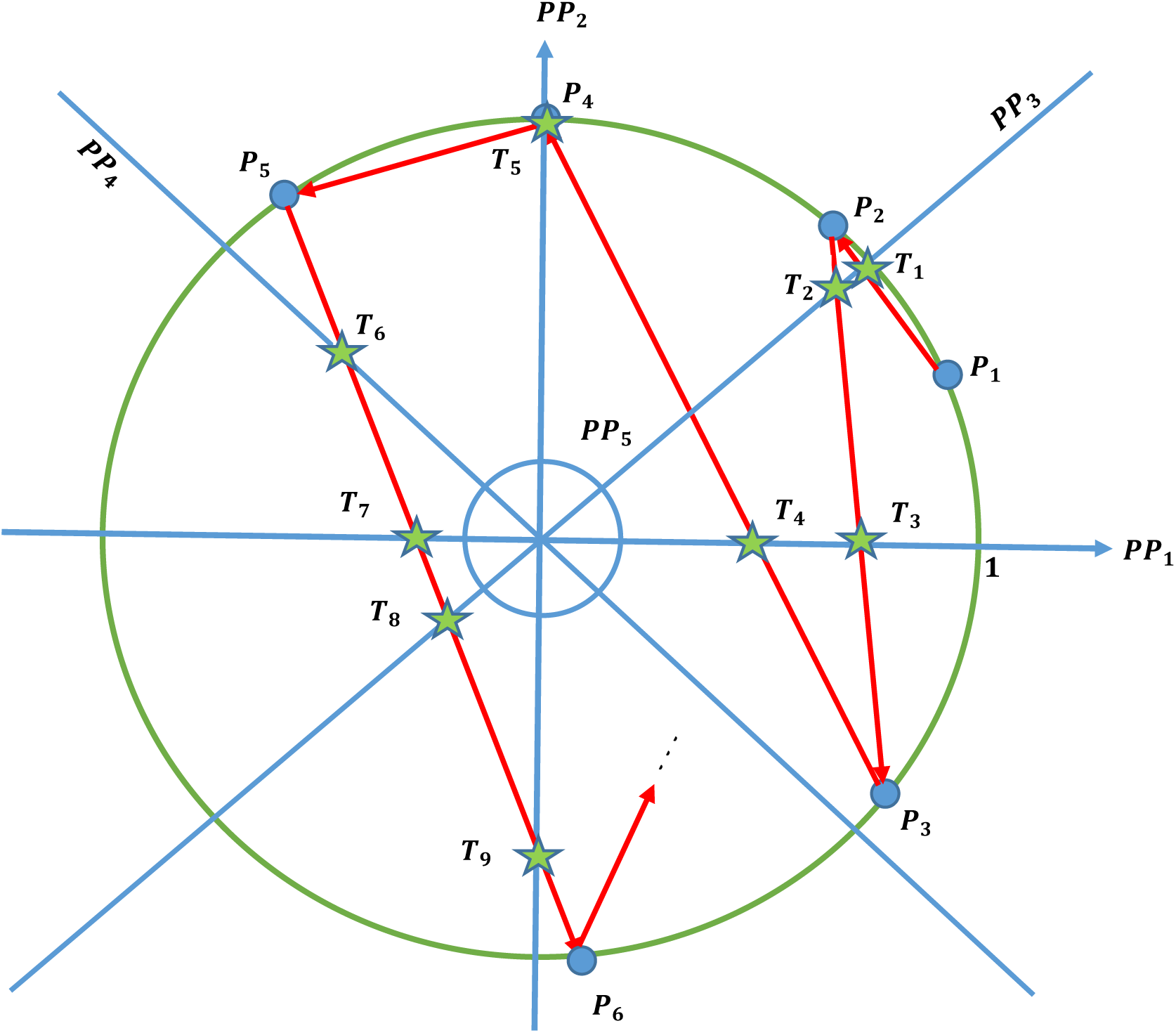
Angle plot (AP) for the hypothetical phase space and Poincare planes. Stars show the intersections of AP with the planes.

As it is obvious, in Fig 5 Poincare sections are defined and *T*_*i*_ points show the intersections (or Poincare points). Table 2 represents Poincare planes and the abbreviations.

**Table 1.** Poincare planes used in this study

| # | Abbreviation | Description |
| --- | --- | --- |
| 1 | PP1 | X axis |
| 2 | PP2 | Y axis |
| 3 | PP3 | Diagonal line (first and third quadrant bisector) |
| 4 | PP4 | Perpendicular to diagonal line (second and fourth quadrant bisector) |
| 5 | PP5 | Circular plane with the radius of $r = 0.001$ |

**Table 2.** Extracted features from AP and PPs for source identification

| # | Feature Description | Abbreviation |
| --- | --- | --- |
| 1 | Average of angle values | *AveAP* |
| 2 | Variance of angle values | *VaAP* |
| 3 | Skewness of angle values | *SkAP* |
| 4 | Kurtosis of angle values | *KuAP* |
| 5 | Median of angle values | *MeAP* |
| 6 | Shannon's entropy of angle values | *ShAP* |
| 7 | Length of the angle time series | *LeAP* |
| 8 | Number of intersections with PP1 | *NPP<sub>1</sub>* |
| 9 | Number of intersections with PP2 | *NPP<sub>2</sub>* |
| 10 | Number of intersections with PP3 | *NPP<sub>3</sub>* |
| 11 | Number of intersections with PP4 | *NPP<sub>4</sub>* |
| 12 | Number of intersections with PP5 | $NPP_5$ |

As mentioned before, in this study just the angle values are considered and features are extracted based on AP and the proposed Poincare planes. Statistical features containing mean, variance, skewness and kurtosis are extracted from AP. Features employed for source identification are explained in Table 3. Statistical features including average, variance, skewness and kurtosis of the angle values are extracted. Number of Poincare intersections for each PP is also considered as a feature.

**Table 3.** Average and standard deviation of classification accuracy for all datasets

| Length | Classifier | SCEEG | SSCEEG1 | SSCEEG2 | RCEEG |
| --- | --- | --- | --- | --- | --- |
| 10 s | MLP | $75.72 \pm 6.55$ | $76.64 \pm 5.52$ | $76.16 \pm 6.33$ | $75.62 \pm 6.51$ |
| | KNN (K=20) | $76.53 \pm 7.41$ | $76.17 \pm 6.09$ | $76.32 \pm 7.11$ | $75.85 \pm 7.16$ |
| | Bayes | $77.04 \pm 8.69$ | $76.39 \pm 5.36$ | $76.22 \pm 6.27$ | $75.49 \pm 7.23$ |
| | Mixture | $78.12 \pm 5.74$ | $79.06 \pm 5.87$ | $78.93 \pm 5.85$ | $78.86 \pm 5.95$ |
| 30 s | MLP | $76.49 \pm 7.93$ | $75.84 \pm 6.16$ | $75.33 \pm 7.41$ | $75.37 \pm 7.79$ |
| | KNN (K=20) | $76.31 \pm 7.01$ | $75.92 \pm 6.09$ | $75.87 \pm 6.71$ | $75.41 \pm 8.23$ |
| | Bayes | $76.17 \pm 7.39$ | $75.78 \pm 7.04$ | $75.63 \pm 8.08$ | $75.49 \pm 6.23$ |
| | Mixture | $78.01 \pm 7.68$ | $78.17 \pm 7.98$ | $77.93 \pm 7.15$ | $78.86 \pm 5.95$ |
| 60 s | MLP | $77.06 \pm 6.55$ | $75.64 \pm 6.52$ | $75.16 \pm 6.33$ | $75.62 \pm 7.51$ |
| | KNN (K=20) | $76.53 \pm 6.41$ | $75.17 \pm 5.09$ | $75.32 \pm 5.35$ | $75.85 \pm 6.16$ |
| | Bayes | $76.04 \pm 8.69$ | $75.39 \pm 7.44$ | $75.22 \pm 7.29$ | $75.49 \pm 7.23$ |
| | mixture | $78.63 \pm 7.58$ | $77.63 \pm 6.57$ | $78.06 \pm 6.32$ | $76.93 \pm 6.04$ |

**Table 4.**
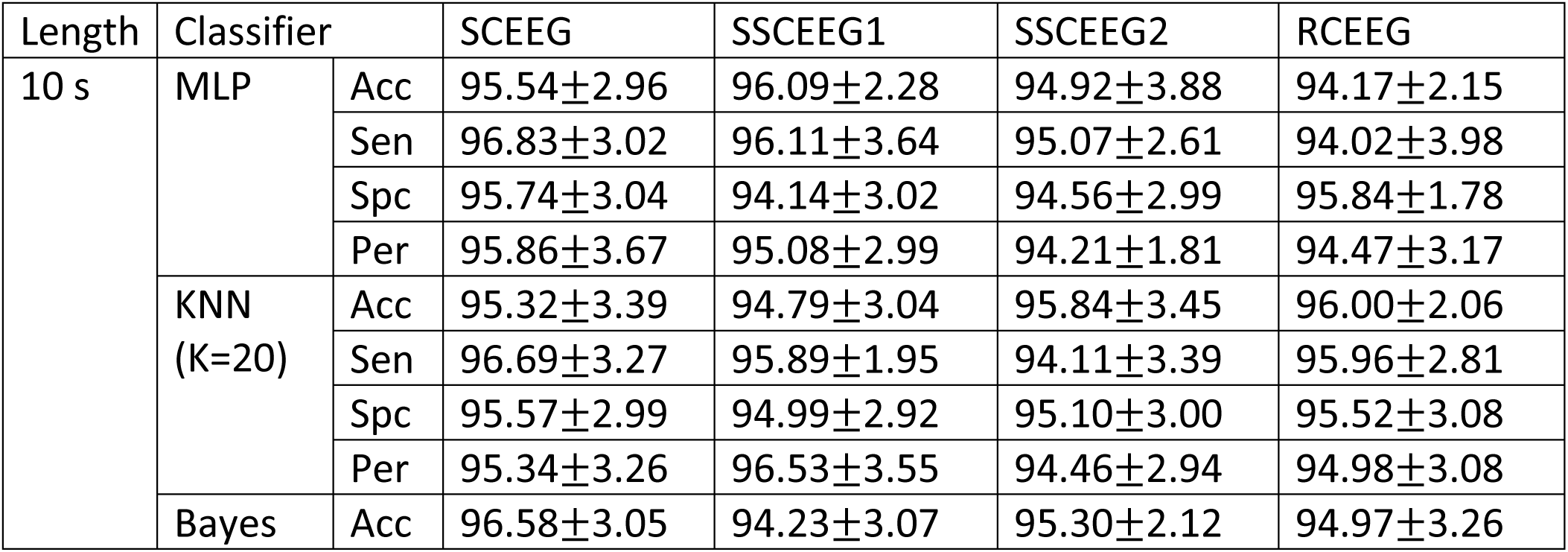
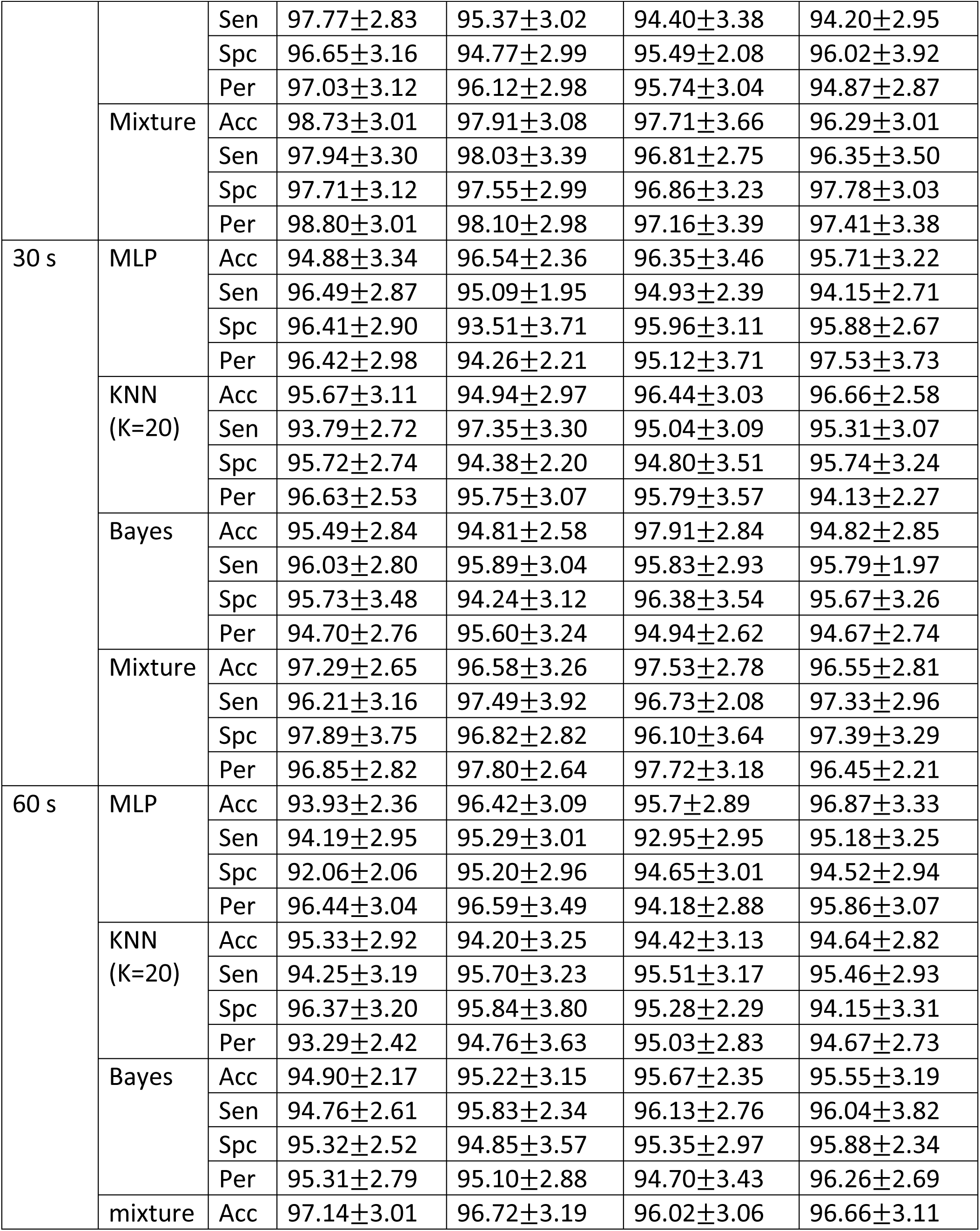
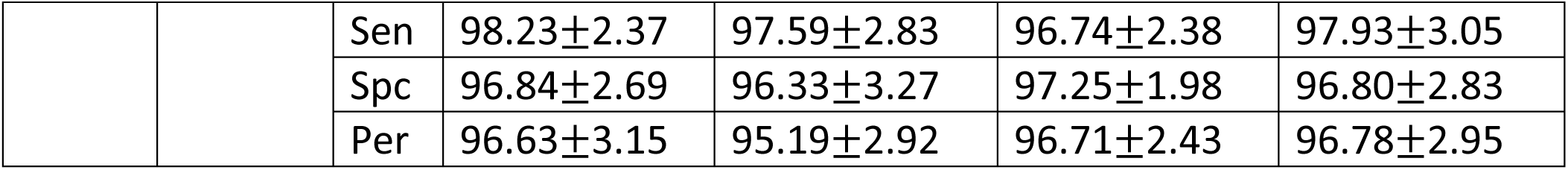
Classification results for recognizing neural and artifactual components

For sake of simplicity just these twelve features are extracted. Results show that all of these features are significant. These features are extracted from each source. Source Identification is performed based on these features. As mentioned before, all features are extracted from AP in order to reduce complexity and processing time.

### 2.5. Classification

K-nearest-neighbor (KNN), Bayes classifier and multi-layer-perceptron (MLP) are three basic and common classifiers which generate immense interest in numerous studies in different fields. KNN, Bayes and MLP are employed in this paper in order to have a more comprehensive study. Due to the wide usage, several pattern recognition toolboxes have implemented these classifiers [REF]. Like other classifiers, these proposed classifiers recognize samples based on two major phases called training and testing steps. KNN is very effective while samples have spherical distribution is the feature space because it classifies samples based on the distances and nearest neighbors. MLP is able to solve complex classification problems. It is also efficient in most machine learning problems since it utilizes error back propagation algorithm to adjust weights and come to generalization in data recognition. We also take advantage of Bayesian classifier’s properties in minimizing the classification error based on probability density functions of training samples [REF]. Since these classifiers have different methods to identify samples, we can compare the results in a more appropriate way. These classifiers are explained precisely in other works like, so we avoid reviewing them here. For more information refer to machine learning and pattern recognition text books like [REF] or articles such as [REF]. Since source recognition is the part of this study, we report classification accuracy for these three basic classifiers in order to compare the results. **\*\*\*BETTER\*\*\***

### 2.6. Wavelet-based artifact removal

Different algorithms can be taken into account to remove artifacts. One can set artifactual components to zero which is not very practical since neural information is very possible to leak into these components. So ignoring all artifactual sources might lead to information loss. Although this approach seems to be very simple, it leads to significant distortion in reconstructed EEGs. On the other hand, a well-known algorithm to suppress artifacts is decomposing artifactual components by wavelet transform. Decomposed sub-bands are denoised by thresholding [10]. Several studies have suggested wavelet with the aim of artifact elimination [54, 59, 60]. The type of wavelet transform varies in each study. It can be discrete wavelet transform (DWT), continuous wavelet transform (CWT) or stationary wavelet transform (SWT) [10]. As it is stated in [54, 61, 62], SWT is superior to DWT and CWT in removing biological artifacts. Additionally, SWT is translation-invariant which suggests its superiority to DWT while removing biological artifacts. According to the results in [54], we employ SWT to denoise detected artifactual components. Fig 6 represents the block diagram of the suggested artifact removal approach using SWT.

**Fig 6.**
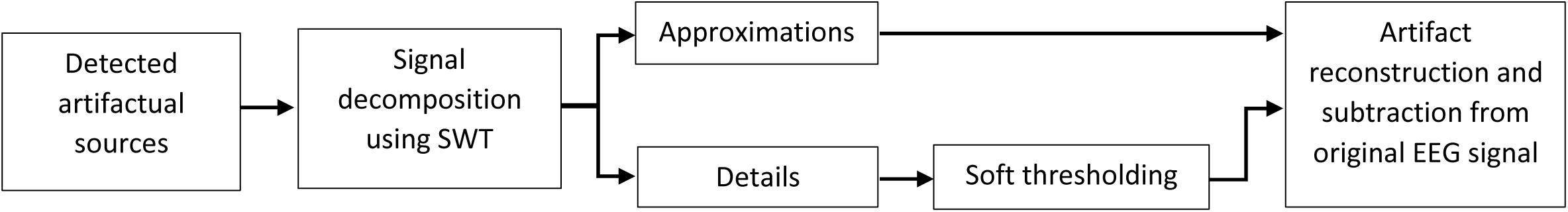
The block diagram for the proposed artifact elimination method based on SWT

**Fig 6.**
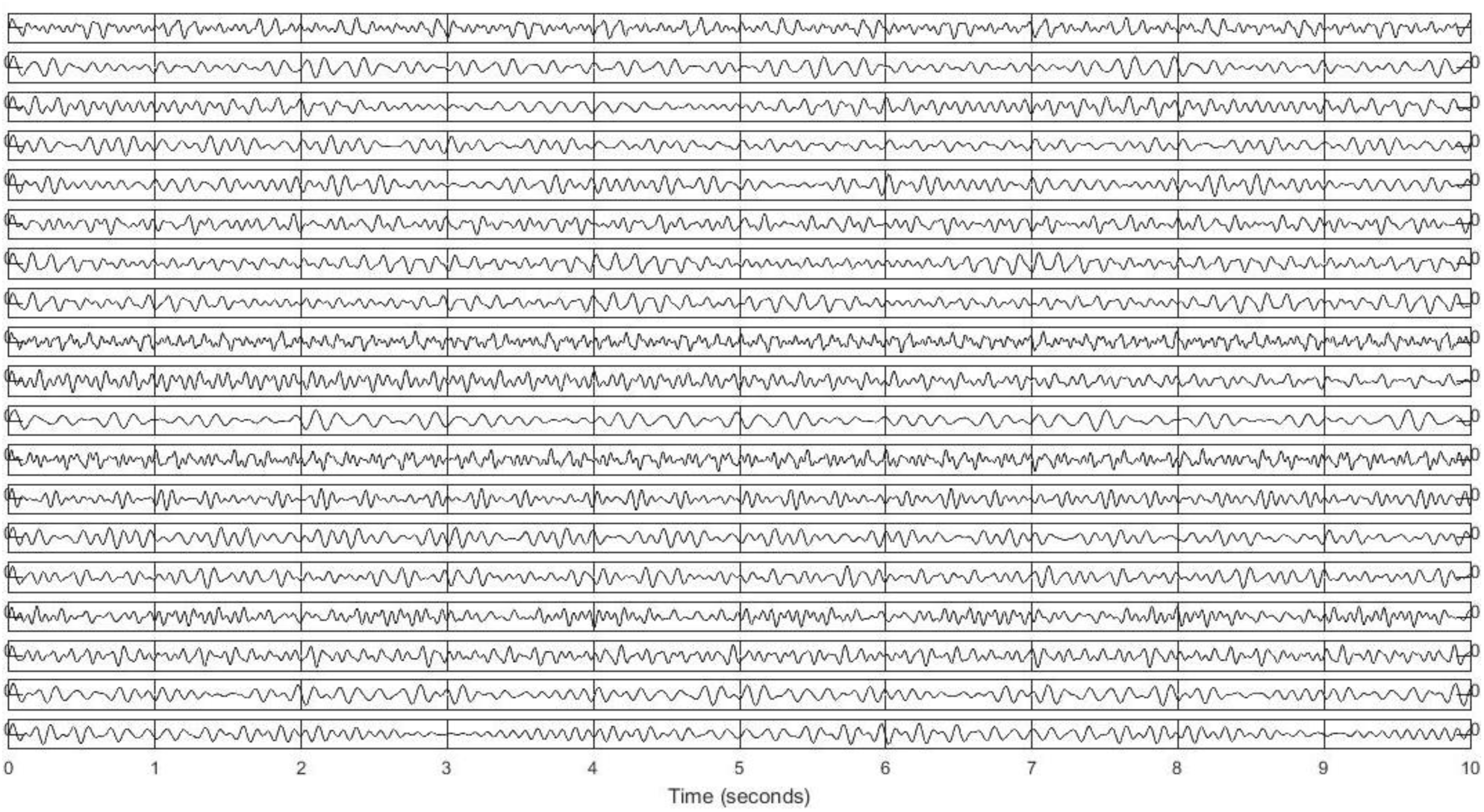
A 10-s illustration of PSEEGs

**Fig 7.**
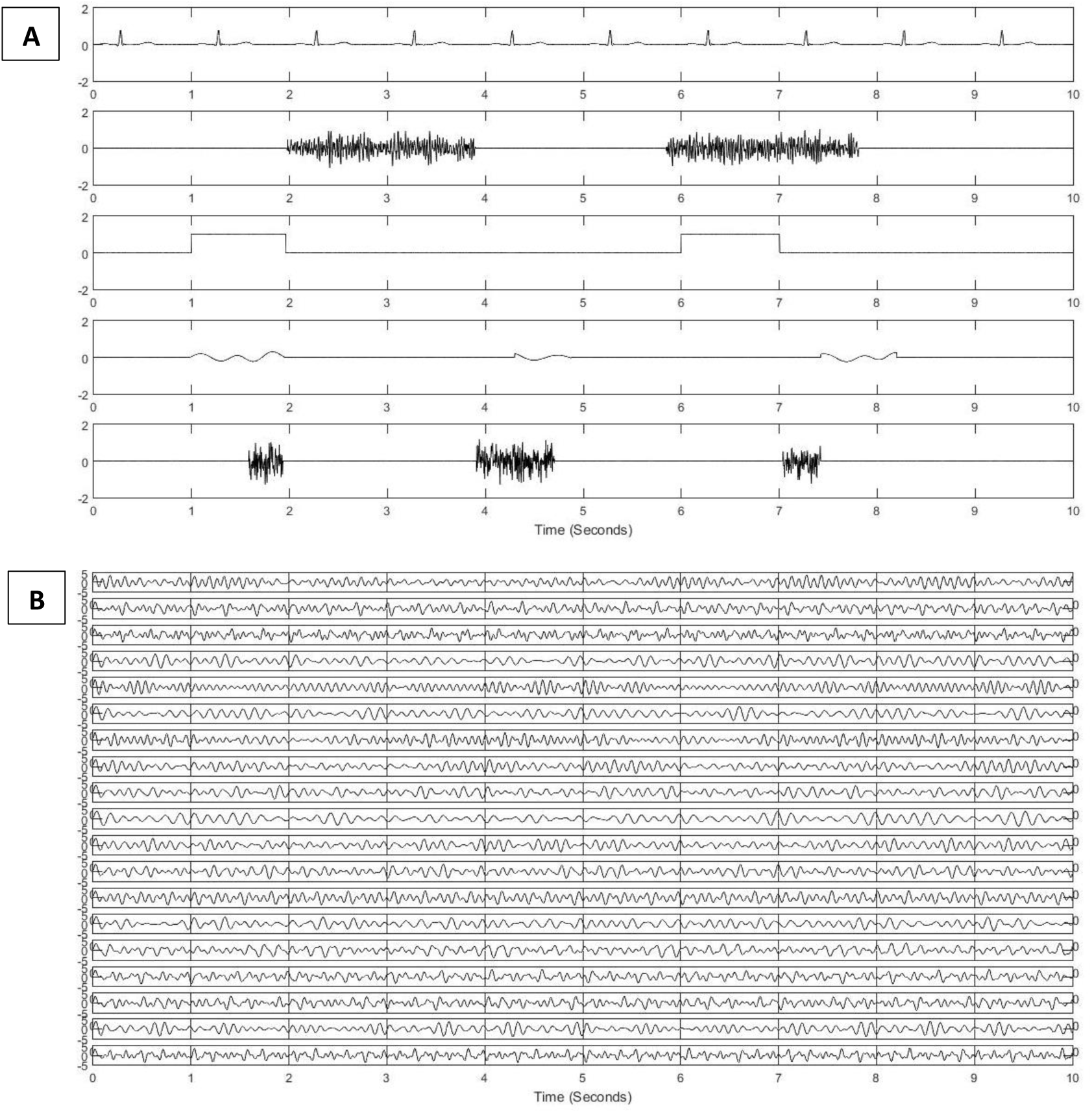
(a) Synthesized artifacts (b) SCEEGs by projecting artifacts at *SNR* = 0.5*db*

We decide to use Haar wavelet because of its advantages in comparison with other wavelet basis functions, five level of decompositions and soft thresholding as it is suggested in [54]. Wavelet analysis results in obtaining approximations and details which correspond to strong artifacts and cerebral information respectively. Artifactual sources are decomposed and sub-bands are taken into thresholding step since in this application approximations correspond to artifacts and obviously details pertain to cerebral activity. So we apply soft thresholding to remove small values in details. Inverse SWT is applied to approximation and thresholded-details to achieve artifacts-only signals. Then the reconstructed artifacts are subtracted from the original signal to have clean EEGs. By thresholding, small values of leaked EEGs would be removed and consequently artifact-only components could be reconstructed, projected back to EEG channels and then subtracted from EEG data. SWT-based approach can be easily implemented using MATLAB wavelet toolbox [63]. The proposed denoising algorithm is quite common, fast and simple. This approach is presented in [54]. Similar to [54], we choose 5 as the level of decomposition and global threshold is computed by MATLAB function *ddencmp*.

### 2.7. Source identification and artifact removal performance measures

Although artifact removal methods are mainly evaluated based on different criteria, the evaluation procedure has been always problematic because there is no universal or general quantitative criterion [10]. Method’s effectiveness can be analyzed through visually inspection by experts which is subjective and not standard or by defining some objective measures which are described below. We consider both subjective and objective metrics in this study. Experts label real and synthesized signals and also extracted sources. So classification performance is the first performance measure. In addition, artifactual sources are suppressed and then “clean” EEG is reconstructed. Therefore, we can define other metrics to evaluate the proposed artifact removal method. Based on the previous studies [REF], some common measures are introduced as the evaluation criteria in this study.

#### 2.7.1. Classification performance

Classification accuracy is defined based on the proportion of the number of correctly classified test samples and the number of total test samples. Considering employing k-fold cross validation in this study, average classification performance (ACP) is calculated and reported for each classifier.

#### 2.7.2. Temporal and Spectral relative root-mean-square error

Artifact removal systems can be evaluated through relative root-mean-square error (RRMSE) in time domain. Several studies consider this factor as an artifact suppression evaluation parameter [50, 57, 58]. RRMSE is defined in time domain as below:

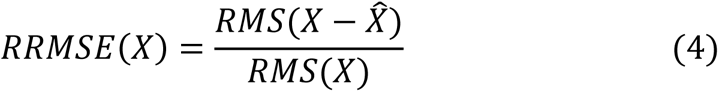

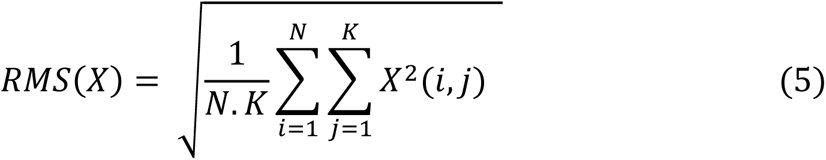

Where *X* and 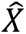 are contaminated (i.e. before artifact removal) and reconstructed (i.e. after artifact removal) EEGs respectively. It can be easily expanded to frequency domain in order to estimate relative root-mean-square error using power spectral density (PSD) which leads to another measure (i.e.*RRMSE*_*PSD*_) described as following:

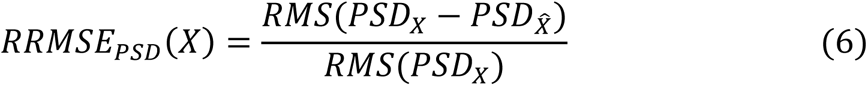

Whit *PSD*_*X*_ and 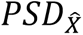 indicating PSD of the clean EEG and denoised EEGs respectively. This measure enables us to analyze the results and evaluate the method with respect to spectral properties of EEGs.

#### 2.7.3. Average correlation coefficient (ACC)

Correlation coefficients (CCs) between original EEGs (not corrupted) and reconstructed ones are useful metrics to evaluate how effectively the proposed artifact removal method can eliminate artifacts. For simulated and semi-simulated signals, original EEGs and artifacts are available. Therefore average correlation coefficient (ACC) could be an evaluation measure. The third criterion in this study is the ACCs over all reconstructed EEG channels with respect to the corresponding original EEG channels [50].

#### 2.7.4. Average mutual information (AMI)

Correlation coefficients cannot fully describe the similarity between two signals or time series. Therefore we decide to employ mutual information (MI) as an index to evaluate the similarity of signal dynamics between original EEGs and reconstructed ones. Average mutual information (AMI) values are computed over all channels as another evaluation parameter. Several studies have applied AMI to quantify their methods [54, ref]. MI is computed in Eq.4 as

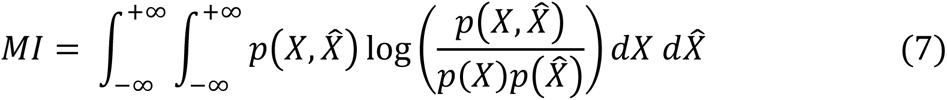

Where 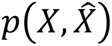 is the joint probability density function of *X* (i.e. original EEG) and 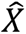 (i.e. reconstructed EEG after artifact removal). *p*(*X*) and 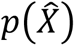 represent marginal probability density functions of *X* and 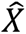 respectively. Since AMI indicates the relevance between two signals, it is clear that the larger the AMI is the more effective the proposed method will be [54]. AMI is employed as another measure of capability of artifact elimination.

### 2.8. Database

#### 2.8.1. Simulated Data

Generating simulated EEGs are introduced in [48] based on the phase-resetting theory. According to [49], EEGs can be reconstructed by adding four sinusoids with randomly chosen frequencies varying from 4 to 30 Hz. Frequency values are selected independently and randomly to synthesize EEGs. So we can easily construct pure-simulated EEGs (PSEEG) through adding four sinusoids. This method is also completely explained in [50]. To reconstruct a 1-min single-channel signal, thirty 2-sec segments are generated and concatenated together. Nineteen channels of EEG and also modeled artifacts are reconstructed in this way. Fig 8 shows one example for PSEEGs.

**Fig 8.**
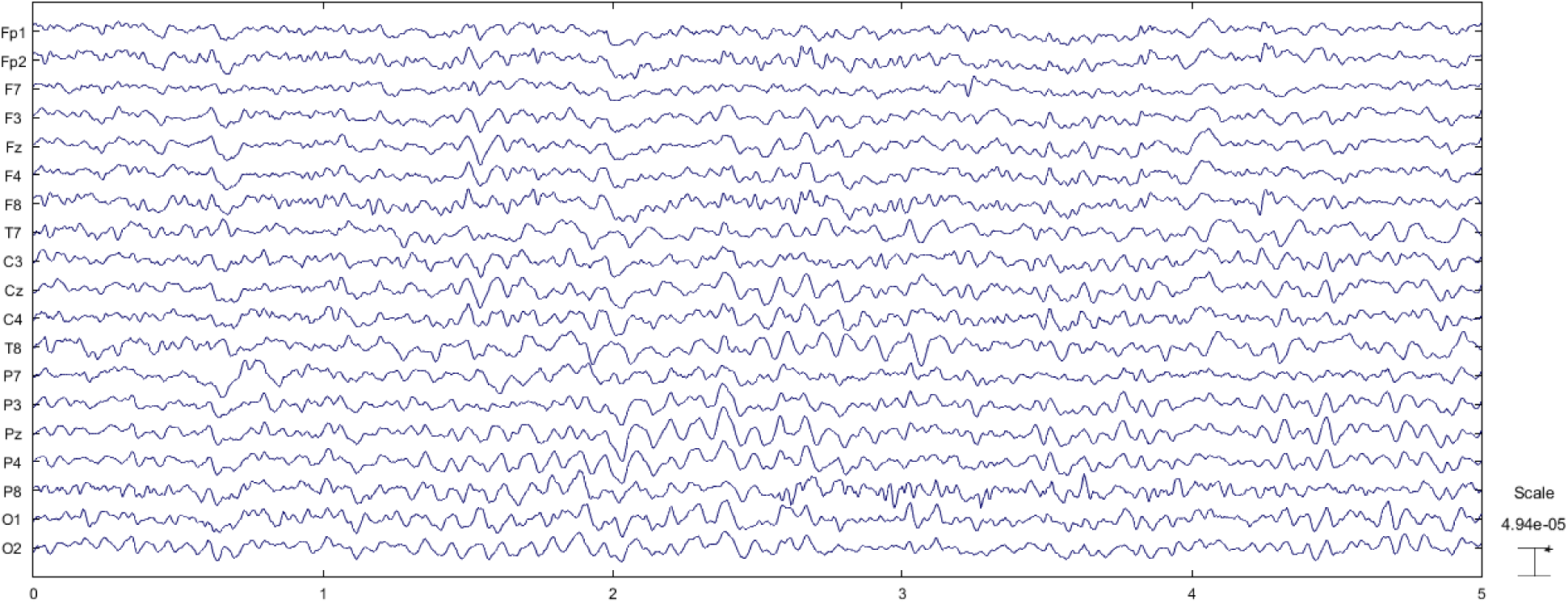
An example for recorded PREEG for 5 s

All signals are recorded or synthesized with a sampling frequency of 256 Hz. We model EEGs and artifacts based on the suggested methods (by previous studies) as bellow:

- EEG: summation of four *sin* functions at random frequencies in the range of 4 to 30 Hz [49],
- ECG: based on previous studies, ECG can be reconstructed by Auto-Regressive (AR) modeling. AR parameters and the order are estimated based on real ECG recordings of participants. Then, artificial ECGs are reconstructed based on AR modeling. We choose AR order as 12 based on Akaike Information Criterion (AIC) and Bayes Information Criterion (BIC) [56] (the average order was 11.6 with the standard deviation of 1.1),
- EMG: temporal muscle activity is modeled by filtering (FIR) random noise in the frequency range of 20 to 60 Hz [53],
- EOG: eye movement is modeled through low frequency square pulses with the frequency of 0.2 Hz [54],
- Eye blinking: we synthesize eye blinking artifact using random noise band-pass filtered between 1 and 3 Hz [53],
- White noise: an unfiltered white noise is employed as an artifact as well.

All five generated artifacts are synthesized in 2-sec segments. We made a decision to generate artifacts in segments with random length varying from 500ms to 2s. In other words, a 2-sec window consists of an artifact based on a random selection. Each modeled artifact is projected to all 19 channels via a random transformation matrix containing at least 10 non-zero random entries and then summed with 19-channel PSEEGs to artificially generate simulated contaminated EEGs (SCEEG). Intensity of artifacts and corresponding channels are randomly selected according to the normal uniform distribution. Based on [50], artifacts can be added to PSEEGs at different levels of signal to noise ratio (SNR). Eq 3 represents the summation of artifacts and EEGs.

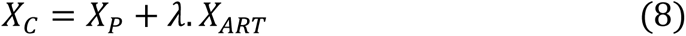

Where *λ* indicates the artifact intensity and totally affects SNR. *X*_*C*_ shows the corrupted 19-channel EEGs. *X*_*P*_ and *X*_*ART*_ demonstrate pure EEGs and modeled 19-channel artifacts respectively. SNR is defined based on Eqs 4 and 5.

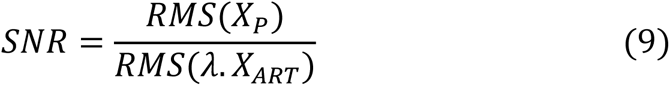

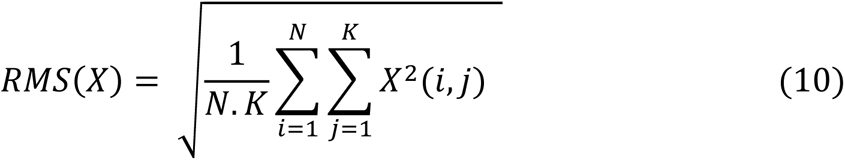

Where *N* is the number of channels and *K* shows time samples. For more information about simulated-contaminated EEG generation refer to [50]. Fig 9 illustrates one example of generated artifacts and CSEEGs.

**Fig 9.**
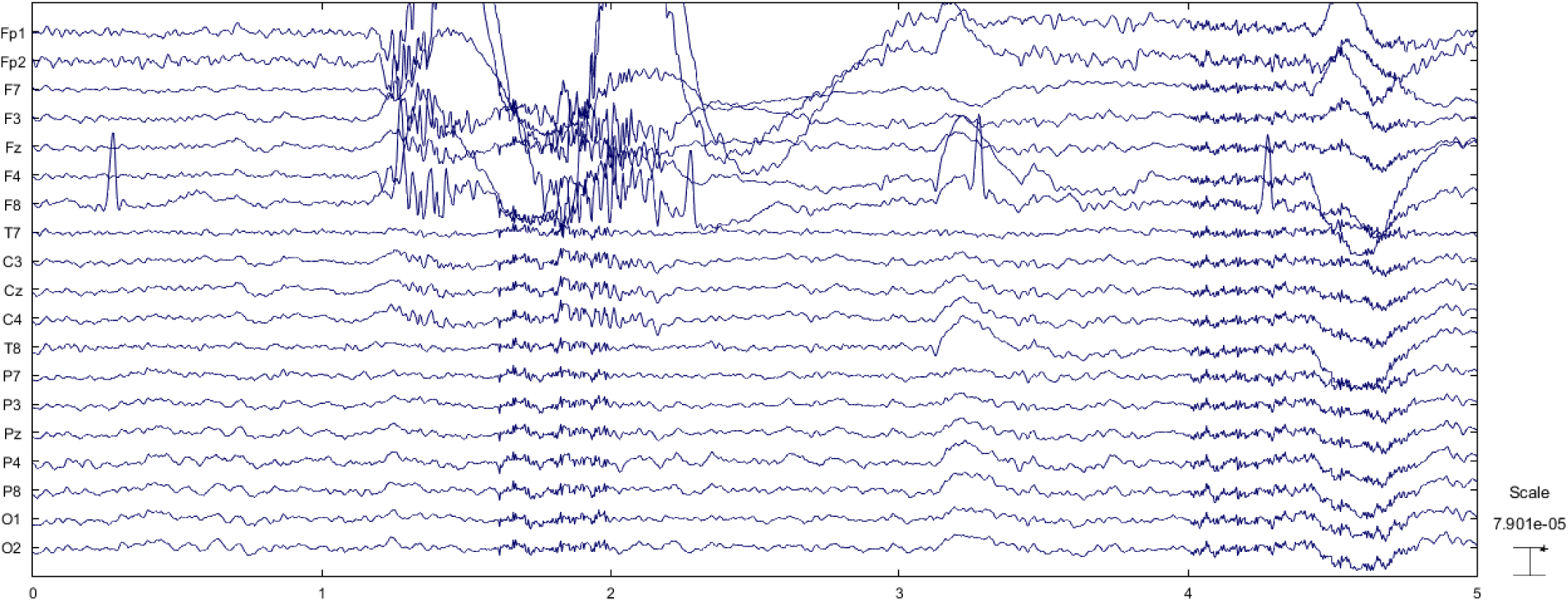
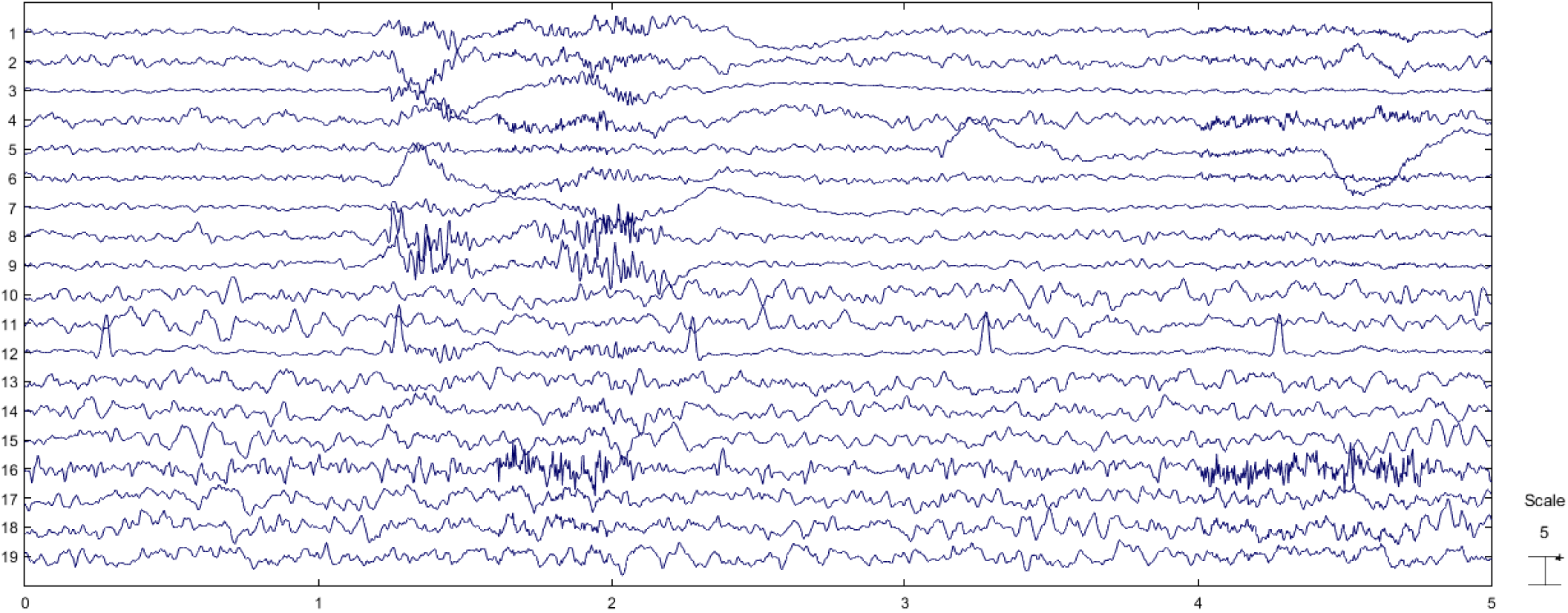
(a) RCEEGs of a subject for 5s (b) extracted sources using SOBI algorithm

#### 2.8.2. Real Data

The EEG signals are recorded from twenty individuals (10 males). 19 Ag/Ag-Cl electrodes according to the 10-20 international standard are placed on each subject’s scalp. EEGs are acquired and sampled at 256 Hz for 1 minute in each trial. Each individual participates in 20 separated trials. 19-channel EEGs are recorded as source identification from fewer electrodes is more challenging and also less time consuming. Furthermore, with 19 electrodes the proposed method is more likely to get employed in other BCI applications. EEGs are recorded while normal subjects are sitting in a comfortable fashion with their eyes open [52]. In the first ten trials for each individual, subjects are acquired not to move their head, jaw or eyebrows. Also eye blinking or movements are visually inspected and not considered in the database. Recorded EEGs are filtered through conventional filtering methods such as bandpass (4-60 Hz) and 50-Hz notch filters based on previous studies like [51, 52] in order to have clean EEGs with no artifacts. Three expert clinicians controlled the recording process and justified clean EEGs. The first ten trials are called pure real EEGs (PREEG). We have PREEGs in 19 channels and ten 1-minute trials for twenty subjects. A sample of PREEGs is represented in Fig 6.

In the next phase, subjects are asked to blink both eyes (without squinting), move their eyes (vertically and horizontally) and eyebrows randomly for 1 minute in each trial. Subjects are left free to blink or move their eyes or eyebrows in their natural manner. Movements are performed in separated and different trials. Eye blinking, eye movement and moving eyebrows are performed in the second ten trials. Subjects are previously informed not to move or tilt their head. Vertical and horizontal EOGs and also ECG are captured in both phases with the aim of helping clinicians while recognizing sources. It should be noted that only 19 contaminated EEG channels are used in further analyses and other signals are recorded due to getting monitored by clinicians. EEGs are filtered by conventional bandpass and notch filters. Subjects are controlled visually while recording signals and movements are recorded in time course. In this phase, individuals participate in ten trials to have real contaminated EEGs (RCEEG). Then RCEEGs and extracted sources via SOBI algorithm are investigated and analyzed by clinicians in order to label sources. Fig 7 shows an illustration of RCEEGs and extracted sources through SOBI algorithm and labeled by experts. Experts are inquired to put each source in one category from all 6 groups containing EEG, ECG, EMG, EOG, eye blink and white noise.

Since artifacts including ECG, eye blinking, EMG, EOG and white noise are very important and prominent in most BCI applications, our focus in this study is on these common artifacts and other artifacts like head movement, power-line noise and electrical shift are ignored [10, 53]. Expert clinicians including three neurophysiologists are informed to control the experiments and label extracted source based on the mentioned artifacts. In this phase we have RCEEGs in 19 labeled sources and ten 1-minute trials for each of twenty subjects.

#### 2.8.3. Semi-simulated Data

To further investigate and study the proposed method, two semi-simulated datasets are provided. In the first dataset, EEGs are taken from PREEGs and artificially-generated artifacts are randomly projected and summed at different *SNR* values. Then extracted sources are identified by experts. PREEGs from each subject are summed with generated artifact described in the previous sub-section. The first set of semi-simulated contaminated EEGs (SSCEEG1) are reconstructed Using Eq. 3 at different *SNR* values. Fig 10 illustrates one typical 5-s SSCEEG at *SNR* = 0.5.

**Fig 10.**
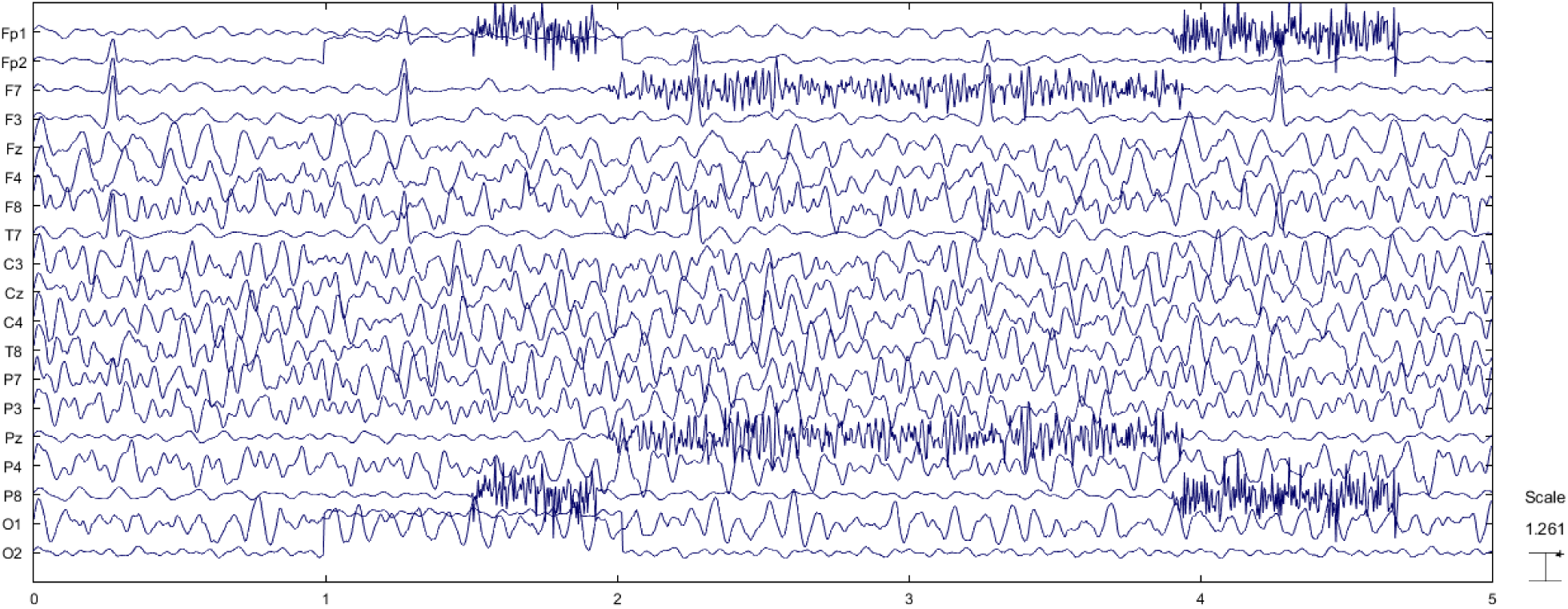
Illustration of 19-channel SSCEEG1s at *SNR* = 0.5

As it can be seen in Fig 10, synthetic artifacts which are explained in the previous sub-section are projected with varying intensities and then added to pure EEG recordings.

For the second semi-simulated dataset, we use EEG signals which are randomly selected from PREEGs. We also record EEGs from 20 other individuals asked to move their eyebrows, blink both eyes and move their eyes horizontally and vertically. Other types of mentioned artifacts like ECG or white noise are seen in the recordings. Individuals are acquired not to move their head. Experts justify artifacts and artifact time courses are visually inspected and recorded. Then artifacts are extracted via FastICA algorithm and identified by expert neurologists. Those extracted artifacts are just considered and then projected back to PREEGs for further analyses. Fig 11 illustrates one example for SSCEEG2. In this approach, we have 200 PREEGs (20 subjects, 10 trials) from the first group of participants and 200 samples which are recorded from the second group of individuals. Extracted artifacts are randomly selected and projected to PREEGs at different *SNR*s to reconstruct the second set of semi-simulated contaminated EEGs (SSCEEG2).

**Fig 11.**
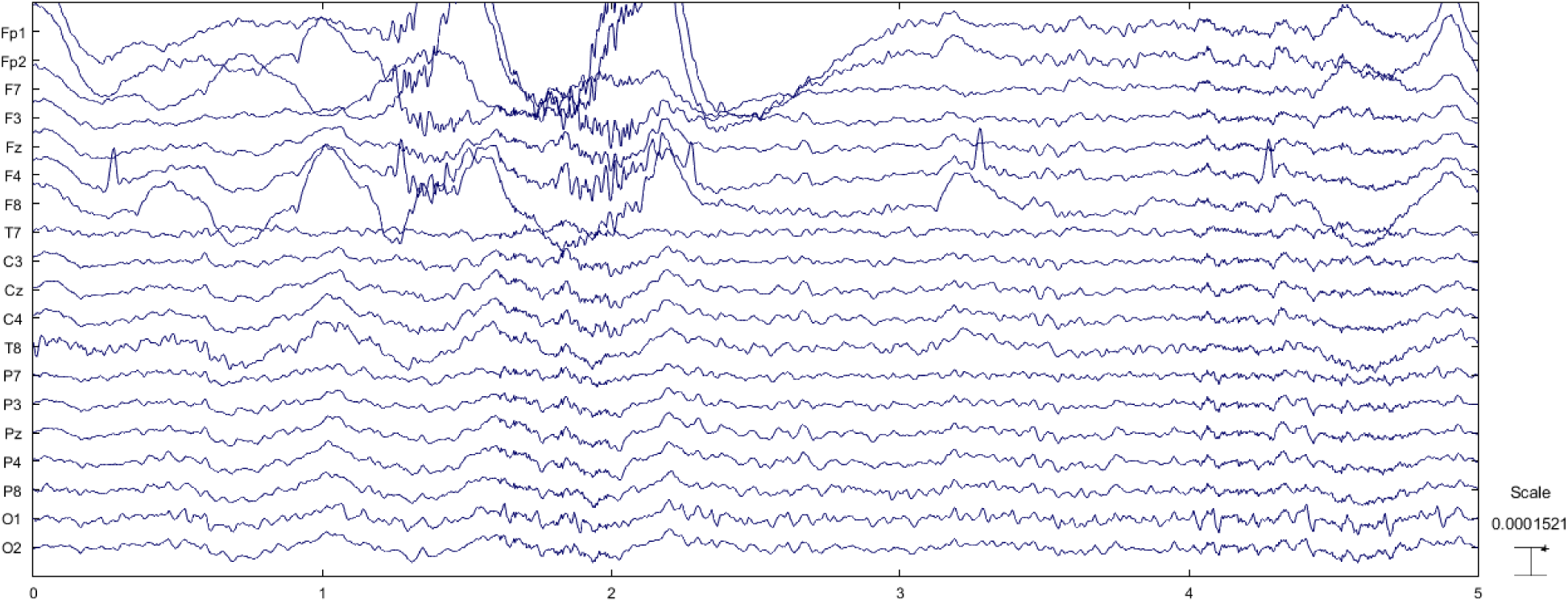
(a) An illustration of SSCEEG2s at *SNR* = 0.5*db*.

In this case, we have the second type of semi-simulated signals containing real artifacts and real EEGs. Figure 11 shows one example for a 5-s semi-simulated contaminated EEG. Needless to say, artifacts can be detected in Fig 11 easily.

## 3. Results

As mentioned before, four different datasets (SCEEG, RCEEG, SSCEEG1 and SSCEEG2) are provided in this study. 200 different 19-channel simulations or recordings are considered for each dataset. For each dataset, contaminated EEGs are fed into SOBI algorithm which is claimed to be a very effective method in separating biological signals and artifacts (specially in the case of EEG) [10]. We assume that the number of sources is equal to the number of channels (*M* = *N* = 19). All extracted sources are analyzed and labeled by experts. Generated or recorded signals and also extracted sources are investigated and monitored by expert neurologists. Samples about which reviewers do not have the same opinions are ignored and removed from the datasets. Experts label all extracted sources and control the reconstructed EEGs. Estimated sources are reconstructed in angle space and mentioned features are extracted based on AP. Mentioned classifiers and also the mixture of the classifiers are employed to classify EEGs and artifacts. We use ten-fold cross validation which means nine folds train the classifiers and one fold is used to test them each time. Classification performance is really important to evaluate the proposed method of automated artifact identification. As it was said before, three common and well-known classifiers are applied in this study. In testing phase, artifactual sources are identified by classifiers and then decomposed by SWT algorithm to perform the artifact removal procedure. Soft thresholding is considered and removing artifacts is carried out based on the suggested algorithm. After artifact identification and removal, denoised EEGs are reconstructed. We apply the proposed method in this study to different signal length to analyze the results more comprehensively. 10, 30 and 60 time windows are considered for signals in this study. Source identification is of a great deal of importance in the proposed method so all extracted sources are labeled in advance and average classification performance (ACP) for testing samples is reported. Table 5 represents the ACP and also standard deviation of classification accuracy for all datasets. ACP shows the average accuracy while classifying samples in each dataset and at different signal lengths.

**Table 5.** Average and standard deviation values for proposed features extracted from each component. P-value for the t-test is also included.

| # | Feature abbreviation | Ave ± std in the brain activity class | Ave ± std in the artifact class | p-value |
| --- | --- | --- | --- | --- |
| 1 | *AveAP* | 37.46±5.17 | 40.02±4.92 | 0.0657 |
| 2 | *VaAP* | 43.81±12.57 | 49.75±10.02 | 0.0789 |
| 3 | *SkAP* | 0.94±0.18 | 0.69±0.39 | 0.0491 |
| 4 | *KuAP* | 3.67±0.94 | 3.01±0.64 | 0.0476 |
| 5 | *MeAP* | 73.31±5.58 | 60.15±6.70 | 0.0581 |
| 6 | *ShAP* | 982312.37±37.93 | 977375.96±42.12 | 0.0464 |
| 7 | *LeAP* | 3217.03±10.56 | 3102.66±9.36 | 0.0812 |
| 8 | *NPP<sub>1</sub>* | 312.87±8.56 | 198.56±9.03 | 0.0299 |
| 9 | $NPP_2$ | $367.35 \pm 10.11$ | $245.73 \pm 9.26$ | 0.0438 |
| 10 | $NPP_3$ | $312.48 \pm 11.78$ | $456.87 \pm 10.49$ | 0.0327 |
| 11 | $NPP_4$ | $327.87 \pm 12.07$ | $423.87 \pm 9.06$ | 0.0492 |
| 12 | $NPP_5$ | $189.58 \pm 7.92$ | $163.44 \pm 6.65$ | 0.0413 |

Classification results suggest that the proposed features and classifiers are effective enough to identify artifacts and EEGs. It should be noted that six groups of samples containing EEG, EMG, ECG, EOG, eye blinking and white noise are considered in classification. As one can see, all accuracy results are quite high and in the same range. It shows that the signal length is not objective in the proposed method. That is to say that results for real EEGs are really similar to that of simulated and semi-simulated ones. This motivates us to examine the mixture of these classifiers utilizing voting of classifiers. It can be easily seen that the mixture of three classifiers outperforms each of them. Therefore, we apply the mixture of classifiers in further analyses to evaluate the proposed method. Additionally, we can evaluate the proposed method considering the artifact removal aspect. For simplicity, results are given in three following subsections to make better comparison. We bring the results just for 10s EEGs in the following sections for sake of space. Since in most studies in this field it is of a great deal of importance to classify artifacts and brain activities, we also decide to classify components into two classes containing neural and artifactual components. Table 5 represents the results. Three afore-mentioned classifiers and the mixture classification model are employed.

In this table Acc, Sen, Spc and Per indicate classification accuracy, sensitivity, specificity and precision respectively. These measures suggest that how successful the classification process is. For more information about these criteria refer to pattern recognition and machine learning text books such as **[REF]**. Results for 2-class recognition show that the proposed method can effectively recognize artifacts. The average values over all datasets and signal length for Acc, Sen, Spc and Per in table 5 is 96.19, 96.38, 95.97 and 95.76 respectively. It should be noted that ten-fold cross validation is performed to evaluate the classification. Classification performance is much higher than 6-class classification step. So we use this classification model for further analysis. Detected components as artifacts are fed into SWT-based artifact removal in the next step. Taking a close look at Acc and Sen measures, it is evident that the mixture of the classifiers is more efficient and successful in comparison with each of classifiers alone. So extracted sources are classified using the mixture model into two classes and artifactual ones are denoised by the proposed method employing SWT.

To analyze the results more completely, we decide to bring features’ average and standard deviation in table 6. So all artifact components in all datasets are put in one group and sources related to brain activity in the other group. It is worth mentioning that for this analysis all components are normalized to the range of [-1 1] before features’ statistics estimation in order to have the same amplitude range for all components. Average and standard deviation values are computed for all EEG recordings and simulations over each extracted feature. T-test analysis is also carried out to investigate the level of significance for each proposed feature. Most significant features are highlighted. Considering the results in table 6, we can easily find out that most proposed features are significant. As it is clear, all features related to Poincare planes have p-value less than 0.05. It shows the importance of nonlinear analysis of signal dynamics. Poincare planes are able to describe the components characteristics. In addition, some statistical features such as skewness and kurtosis seem to be significant. Besides, 2-class classification is carried out over all normalized components. Features whose p-value is less than 0.05 are selected for each component and then classification is carried out. Evaluation is performed using ten-fold cross validation. Table 7 shows the classification performance over all components. It is worth mentioning that all components regardless to their datasets are put in two classes and then classification procedure is performed.

**Table 6.** Classification performance for the normalized components over all samples

| Classifier |  | 10 s | 30 s | 60 s |
| --- | --- | --- | --- | --- |
| MLP | Acc | $97.63 \pm 1.57$ | $96.36 \pm 1.32$ | $96.43 \pm 1.38$ |
| | Sen | $97.22 \pm 1.32$ | $96.27 \pm 1.61$ | $96.46 \pm 1.51$ |
| | Spc | $97.53 \pm 1.38$ | $97.29 \pm 1.12$ | $95.67 \pm 1.75$ |
| | Per | $96.37 \pm 1.24$ | $96.88 \pm 1.26$ | $96.58 \pm 1.36$ |
| | Acc | $96.94 \pm 1.49$ | $97.42 \pm 1.49$ | $96.31 \pm 1.29$ |
| KNN<br>(K=20) | Sen | 97.12±1.38 | 96.37±1.15 | 96.38±1.31 |
| | Spc | 96.28±1.46 | 97.49±1.63 | 96.49±1.68 |
| | Per | 97.07±1.37 | 97.69±1.22 | 95.93±1.58 |
| Bayes | Acc | 97.38±1.31 | 97.85±1.24 | 96.96±1.79 |
| | Sen | 96.21±1.18 | 96.93±1.36 | 95.41±1.43 |
| | Spc | 97.36±1.23 | 97.27±1.33 | 95.57±1.52 |
| | Per | 96.87±1.45 | 97.28±1.47 | 96.32±1.48 |
| Mixture | Acc | 98.26±1.27 | 97.95±1.18 | 98.27±1.37 |
| | Sen | 98.39±1.04 | 98.01±1.09 | 97.76±1.46 |
| | Spc | 97.63±0.95 | 97.98±1.27 | 97.83±1.45 |
| | Per | 98.84±1.01 | 98.21±0.98 | 98.46±1.33 |

Taking a closer look at the recent results in tables 5, 6 and 8, it is evident that classification performance while using the mixture model is quite high and higher than several previous studies such as [52, 53]. Referring to table 5, we can even determine the type of the artifact with the accuracy more than 75% in all cases and datasets. It is noticeable in tables 6 and 8 that all components regardless to the datasets are classified into two classes with the accuracy and sensitivity more than 96% and 95% respectively. These values can give us a rough estimation of the proposed method being effective and efficient. More results are provided in the following subsections.

**Table 8.** Average processing time at signal length 10 s, 30 s and 60 s over all components

| Signal length | 10 s | 30 s | 60 s |
| --- | --- | --- | --- |
| Processing time | $0.08 \pm 0.02$ | $0.17 \pm 0.03$ | $0.23 \pm 0.05$ |

### 3.1. Simulated data results

Figure 8 shows the simulation results for synthesized EEGs. In Fig. 12(a) artificial and contaminated EEGs are represented in 19 channels. Artifacts can be easily seen in this figure. Some of them are pointed out by arrows. Simulated artifacts in this figure correspond to the mentioned artifacts in Fig. 12(a). Fig. 12(b) demonstrates extracted sources via SOBI algorithm. It is evident that first three sources and the last one pertain to the artifacts. These sources are identified as artifacts using the mixture of classifiers. As it can be easily seen, noticeable EEG is leaked to these sources which is so important. Artifactual sources are taken into SWT-based artifact removal algorithm in order to get artifacts eliminated. Fig. 12(c) shows the output sources of SWT. As it is clear, SWT is effective in artifact removal. Then, EEG channels are reconstructed using the inverse of the mixing matrix. Fig. 8(d) represents the final reconstructed EEGs. Considering the pure EEGs and the results of the proposed method, no appreciable artifact is notified in the results. Moreover, reconstructed EEGs are justified by experts in order to evaluate the results visually. As it can be seen in Fig 12(b), brain activity leaks into artifactual components which is why we should not set these components to zero. These components are decomposed and details are thresholded using SWT.

**Fig 12.**
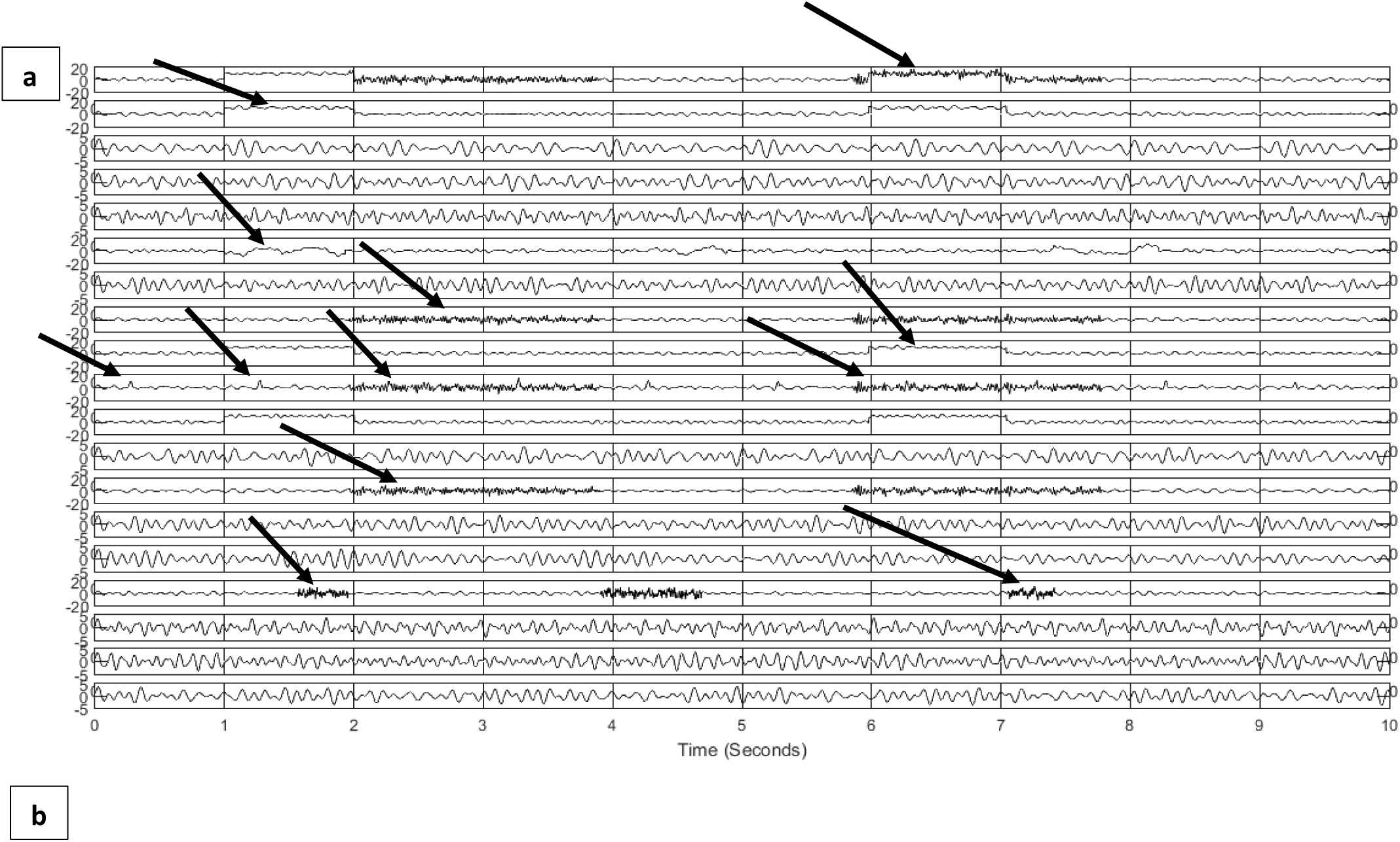
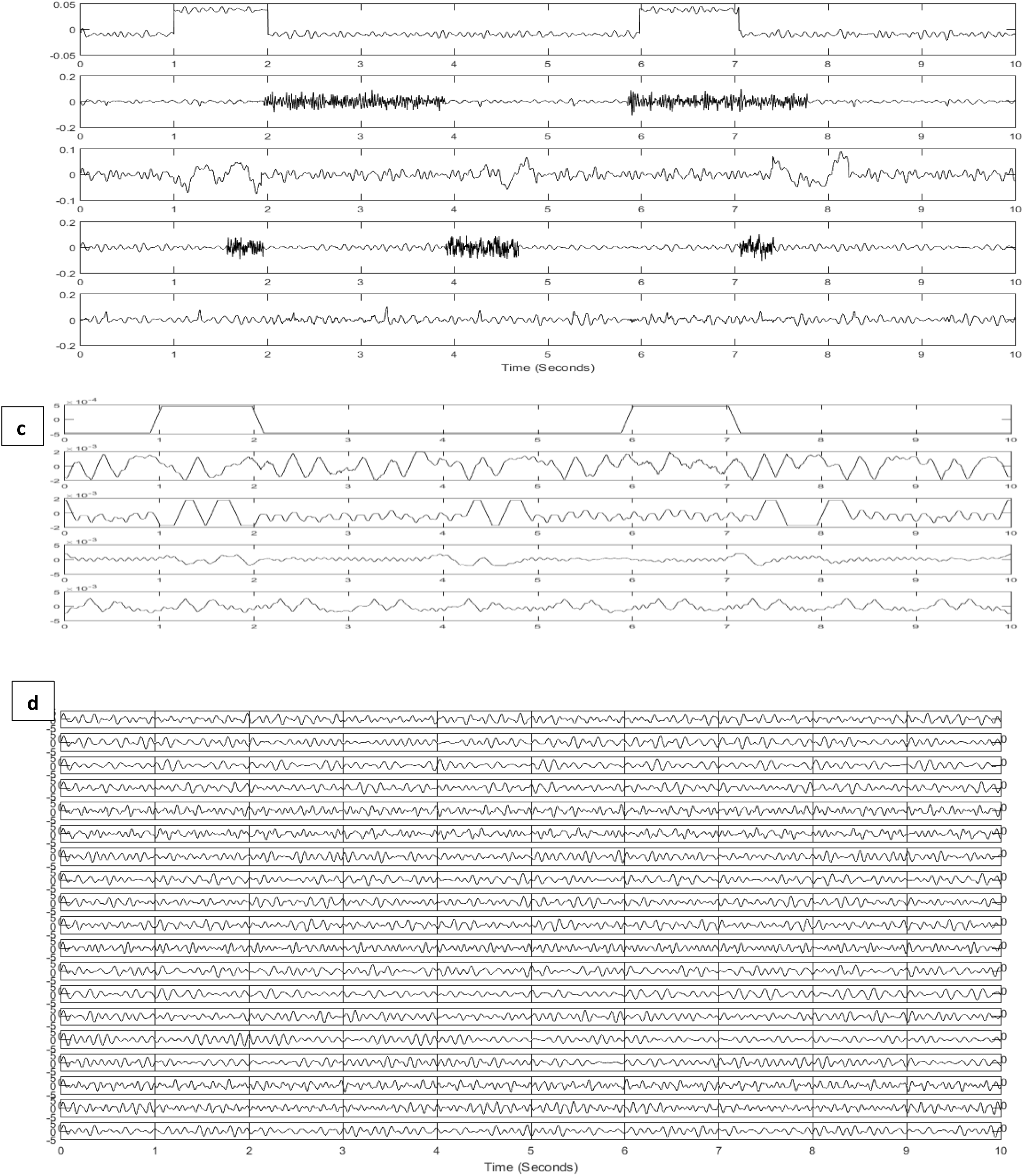
Results of the proposed method for simulated signals (a) simulated and contaminated EEG (b) detected artifactual components (c) denoised components using SWT (d) reconstructed EEG

**Fig 13.**
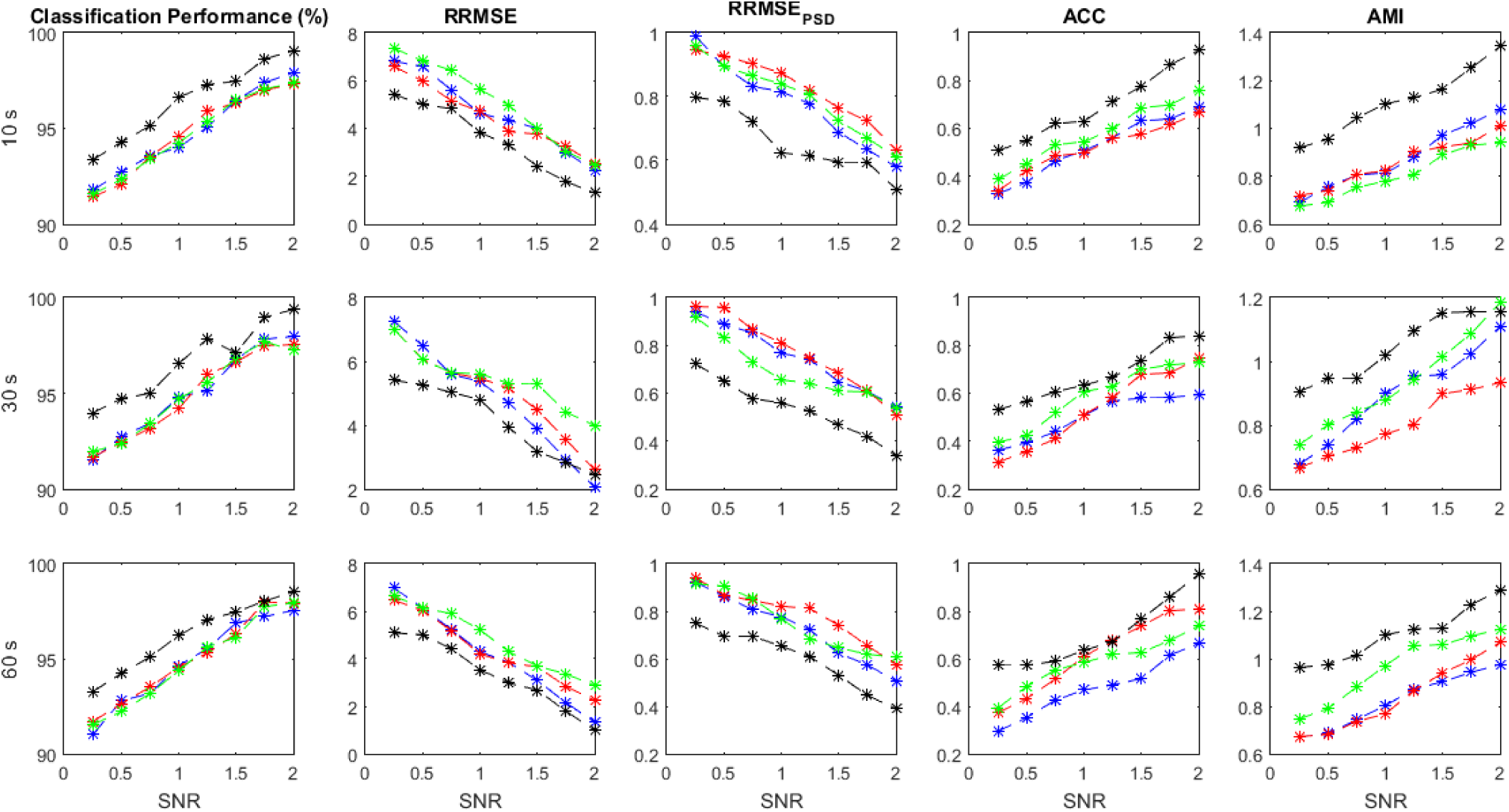
Performance parameters using simulated data at various SNR values in terms of *ACP*, *RRMSE*, *RRMSE*_*PSD*_, *ACC* and *AMI*

In this section, we evaluate the proposed method through 200 independent realizations. In each realization, artifacts are separately generated at random intensities and then added to the simulated EEGs. Contaminated EEGs are taken into SOBI algorithm to get the sources separated. Estimated EEG sources are classified by the mentioned classifiers with respect to extracted features from the AP. Experts monitor and label the sources in advance to ensure the proposed method to work. Artifactual sources are detected and fed into SWT denoising procedure. Artifacts are estimated and then substracted from the original signals in order to achieve artifact-free EEGs. The implementation is carried out at different SNR values to evaluate the method more precisely. *ACP*, *RRMSE*, *RRMSE*_*PSD*_, *ACC* and *AMI* are calculated for each implementation and shown in figure 9. We can compare the effectiveness of different classifiers trained with the suggested features. The green, blue, red and black lines indicate the results for MLP, Bayes, KNN and the mixture of classifiers respectively. Average values for all performance measures are displayed for different length of signals and at different SNRs.

As it is clear the mixture of classifiers outperforms the other classification models at all SNR values. All performance criteria are almost close for MLP, KNN and Bayes but the mixture classification model is more effective. It is worth mentioning that the mixture of classifiers can preserve the original EEGs when they are highly contaminated (e.g. SNR<0.5). It should be noted that average values for each performance parameter at different SNRs are reported and displayed. It should be mentioned that as SNR decreases all performance criteria degrade sharply.

### 3.2. Semi-simulation results

As mentioned before, we have collected two different datasets for semi-simulated signals to investigate the method performance more precisely. The proposed method is applied to both semi-simulated datasets. Fig 14 illustrates the results related to SSCEEG1. This dataset consists of actual pure EEGs contaminated by simulated artifacts. In Fig 14, all performance criteria suggest that the proposed method is quite effective. Similarly, Fig 14 indicates that the mixture of classifiers leads to improved results for different signal length.

**Fig 14.**
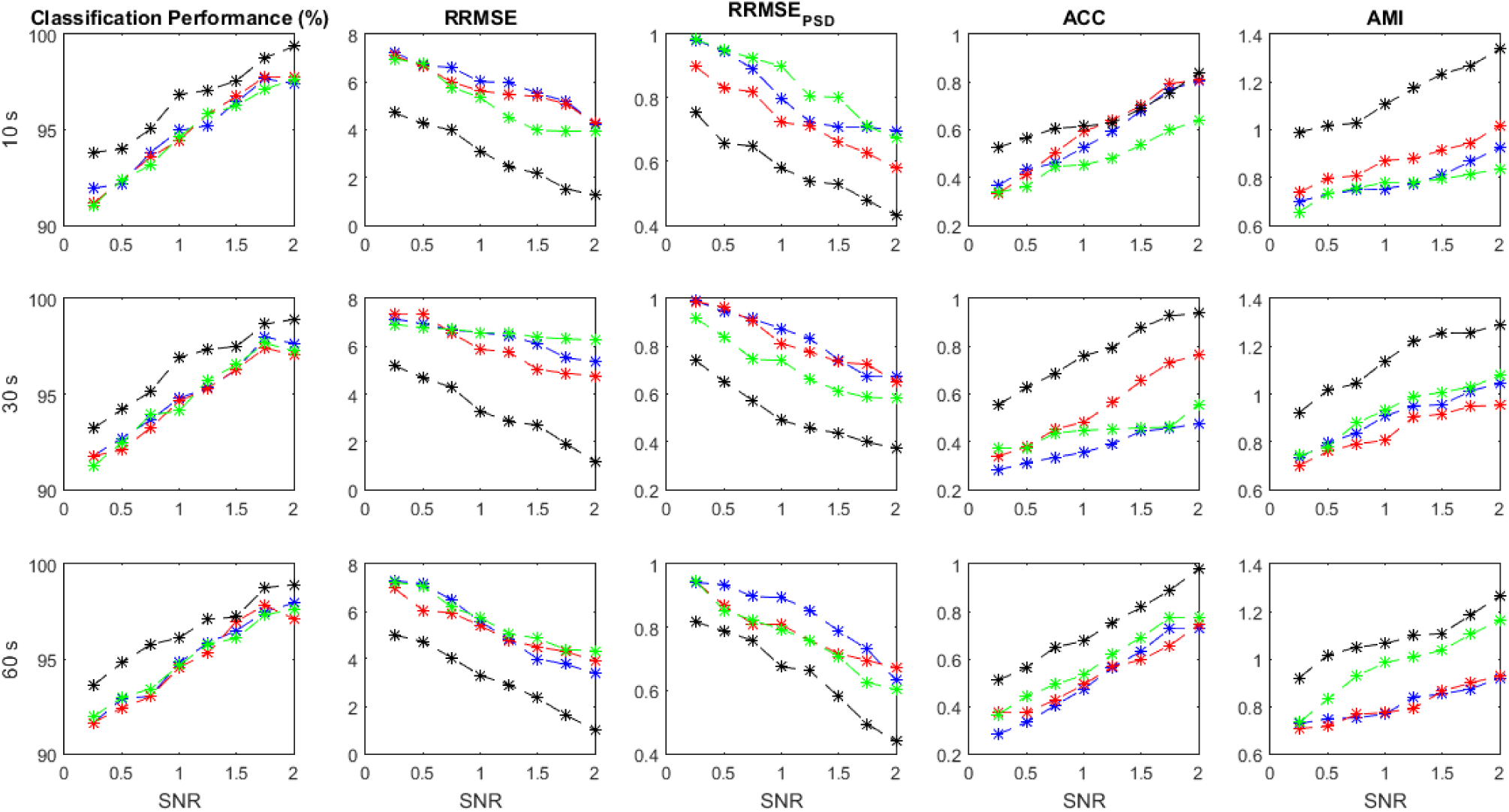
Performance measures for SSCEEG1

Since experts control the generated signals and label all of the extracted sources we can easily measure the performance parameters. Additionally, we have the pure EEGs in both semi-simulated datasets. So evaluation measures containing ACP, RRMSE, RRMSE PSD, ACC and AMI can be calculated for SSCEEG1 and SSCEEG2. Figure 15 shows the performance measures for the second dataset of semi-simulated EEGs called SSCEEG2 at different SNRs. Results for SSCEEG2 are quite similar to that of SSCEEG1. This similarity indicates that the considered EEG model to generated EEGs for simulated dataset is quite reliable and realistic.

**Fig 15.**
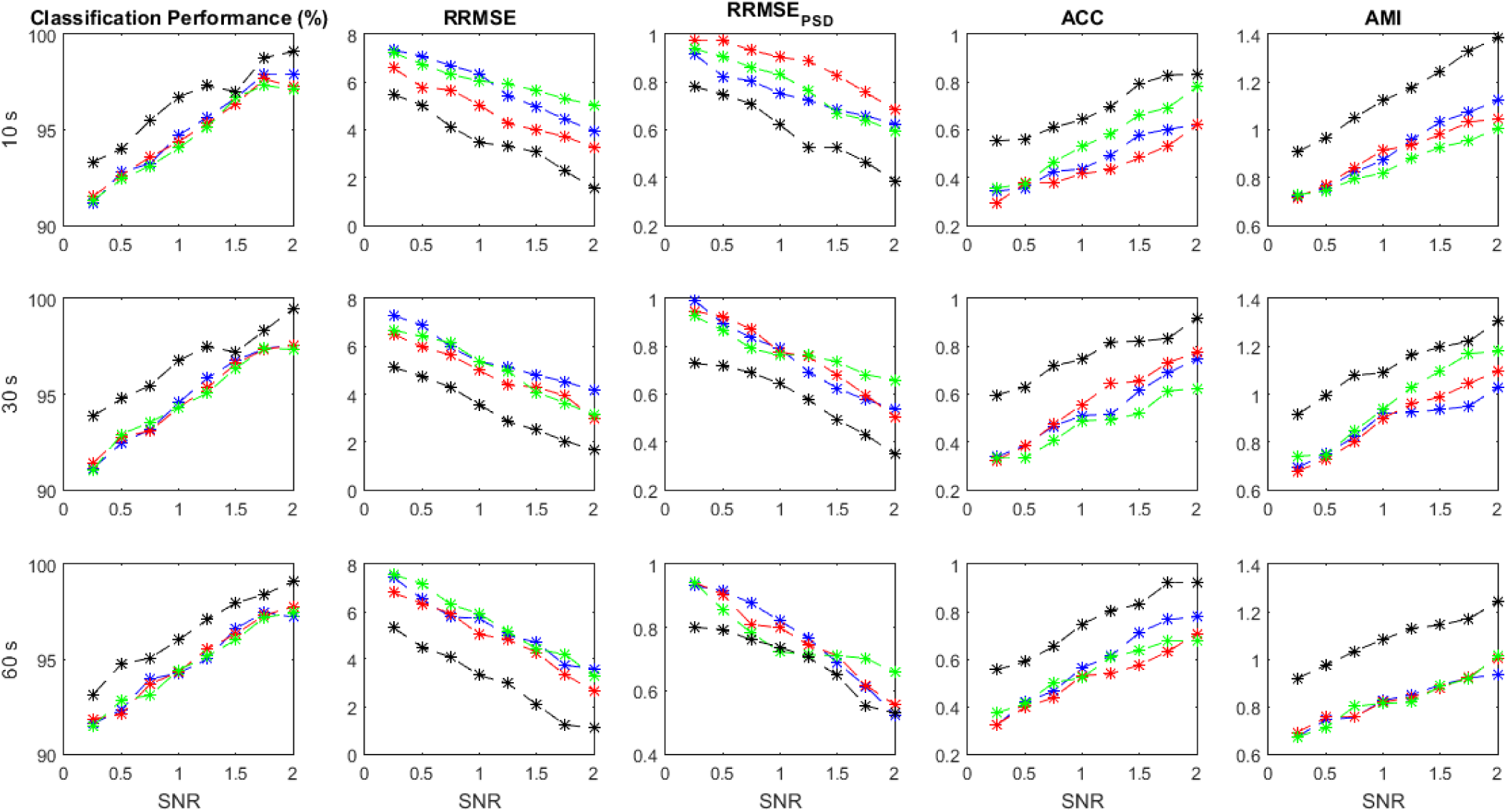
Performance measures for SSCEEG2

Like what was expected, results prove of the proposed model for pure EEGs and also artifacts. Since the results for SSCEEG2 are almost close to that of SSCEEG1, we can conclude that the proposed method is quite practical in real applications and situations. It should be noted that in SSCEEG2, pure real EEGs are randomly selected among participants and randomly-selected real artifacts are added to them at different SNRs. In contrast to SSCEEG2, SSCEEG1 dataset contains pure real EEGs contaminated by artificially-generated and randomly-selected artifacts.

### 3.3. Real data results

In this section we apply the proposed method to real contaminated EEGs. There are 200 EEG recordings and all EEGs and extracted sources are monitored and labeled by experts. These EEGs contain severe artifacts to evaluate the suggested artifact removal procedure. Since there is no ground truth available for real data, we cannot report performance parameters. In other words, for real EEGs evaluation procedure is performed in a quantitative way including visualization criteria like topography or spectral density and temporal analysis. It should be noted that source classification accuracy for real data is previously reported in table 3. As sources are labeled by experts just classification accuracy is available as a quantitative measure. For real contaminated EEGs visual inspection is performed by experts in order to evaluate the proposed method more truly. Figure 16 shows a real contaminated EEG recording from a participants in a 5-second segment.

**Fig 16.**
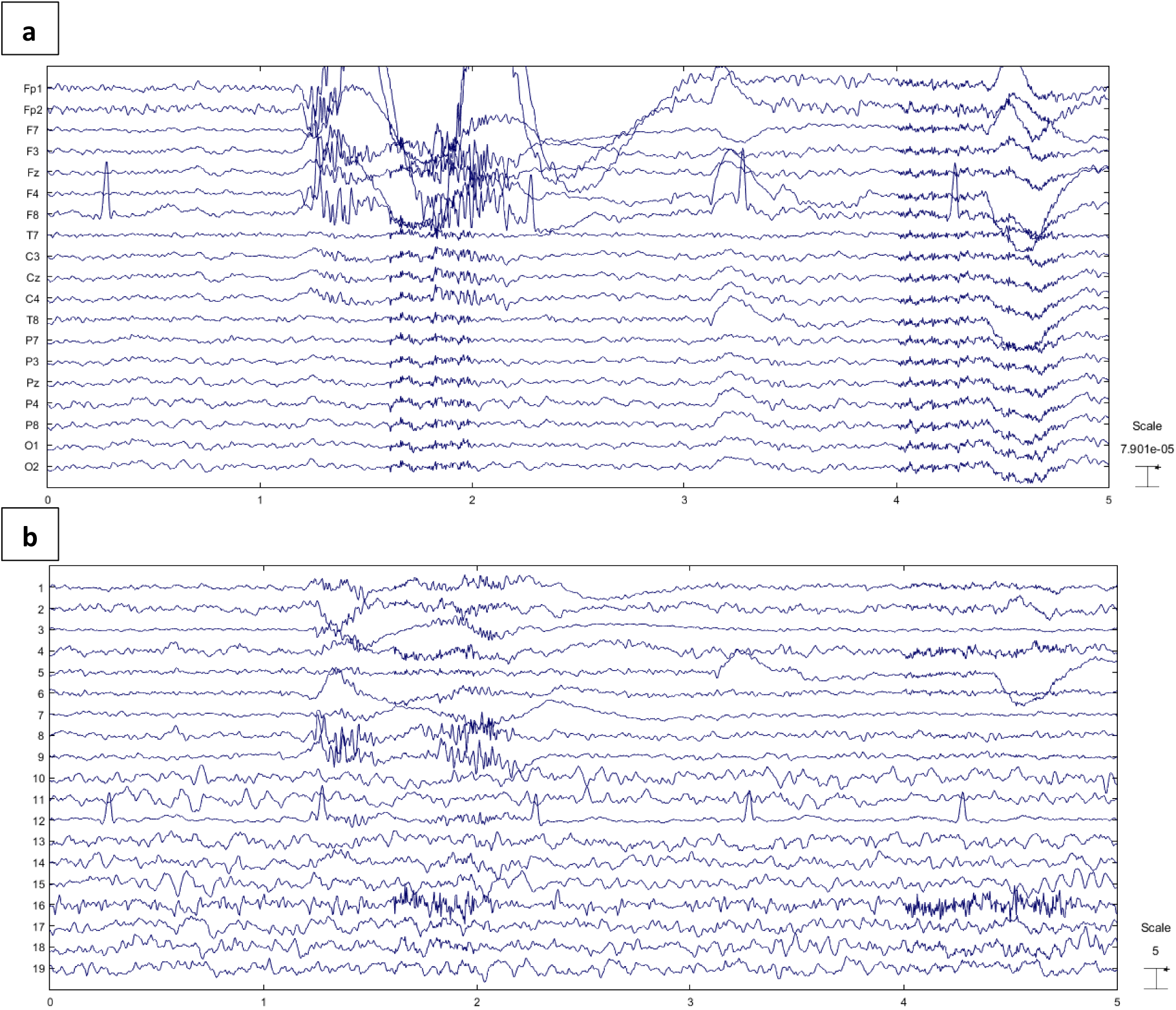
(a) Real contaminated EEG and (b) extracted sources using SOBI

As one can easily see, SOBI is able to separate sources and isolate artifacts. Although the source separation algorithm is quite effective, it is clear that brain activity leaks to most artifactual sources. This motivates us to employ automated artifact detection using the mixture of mentioned classifiers. All detected artifact components are processed via the proposed artifact elimination method based on SWT. In Fig 16, an EEG recording contaminated with all mentioned artifacts is represented. Taking a closer look at extracted sources in Fig 16(b), it is clear that sources 1,2,3,4,5,6,7,8,9,12 and 16 are artifactual. All of these sources are detected by the proposed classification model. Fig 17 shows the reconstructed EEG and its sources after artifact removal.

**Fig 17.**
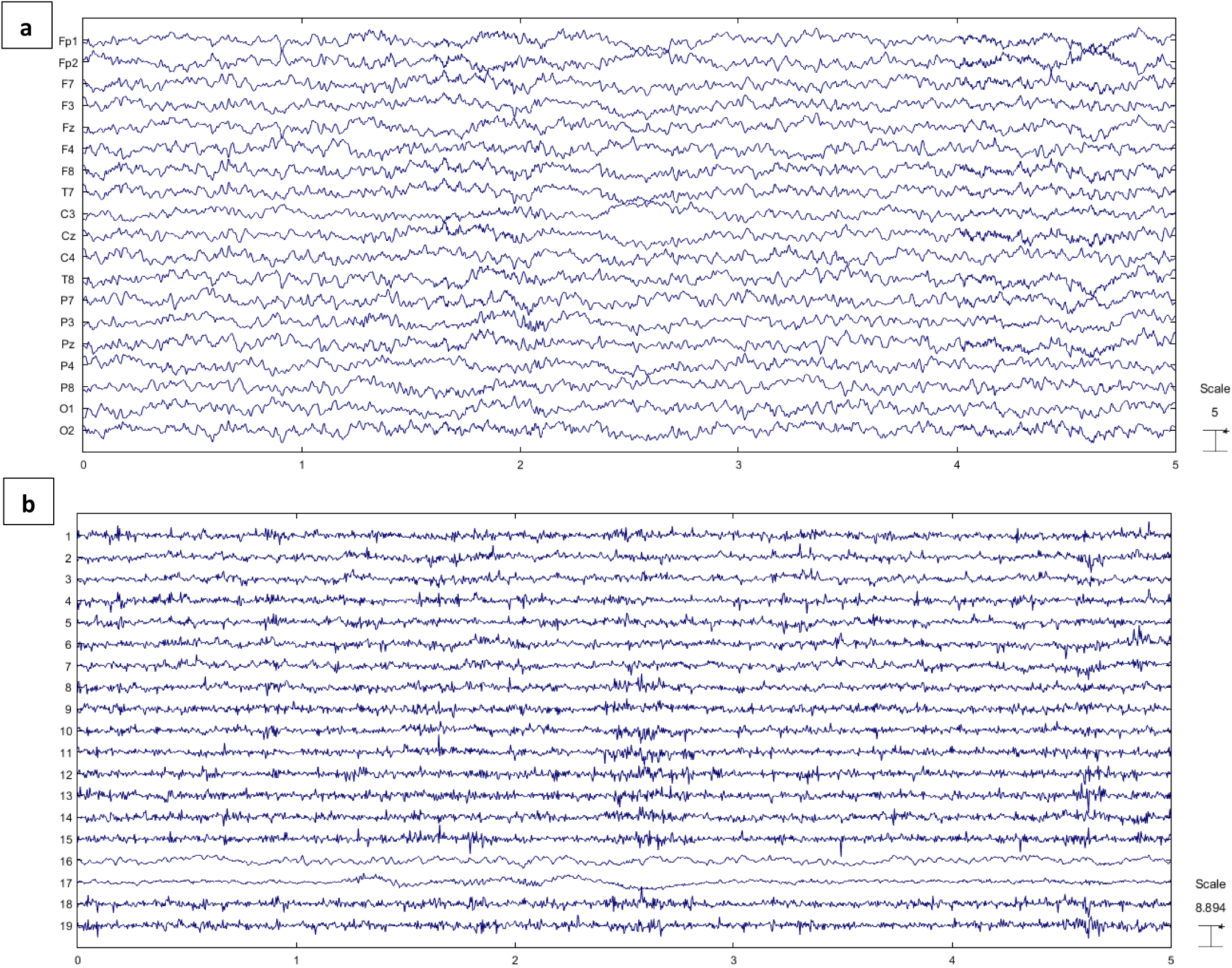
(a) Reconstructed EEG through the proposed method and (b) EEG components.

Some muscle activity can be seen in Fp1, Fp2 and F8. Moreover it might be realized that artifacts such as ones related to eye movement and blinking still remain in the reconstructed EEG. To analyze the results more completely, we decide to consider the topography maps and power spectral density before and after applying the proposed method. Fig 18 illustrates the topography maps for extracted sources and similarly Fig 19 represents power spectral density for channels.

**Fig 18.**
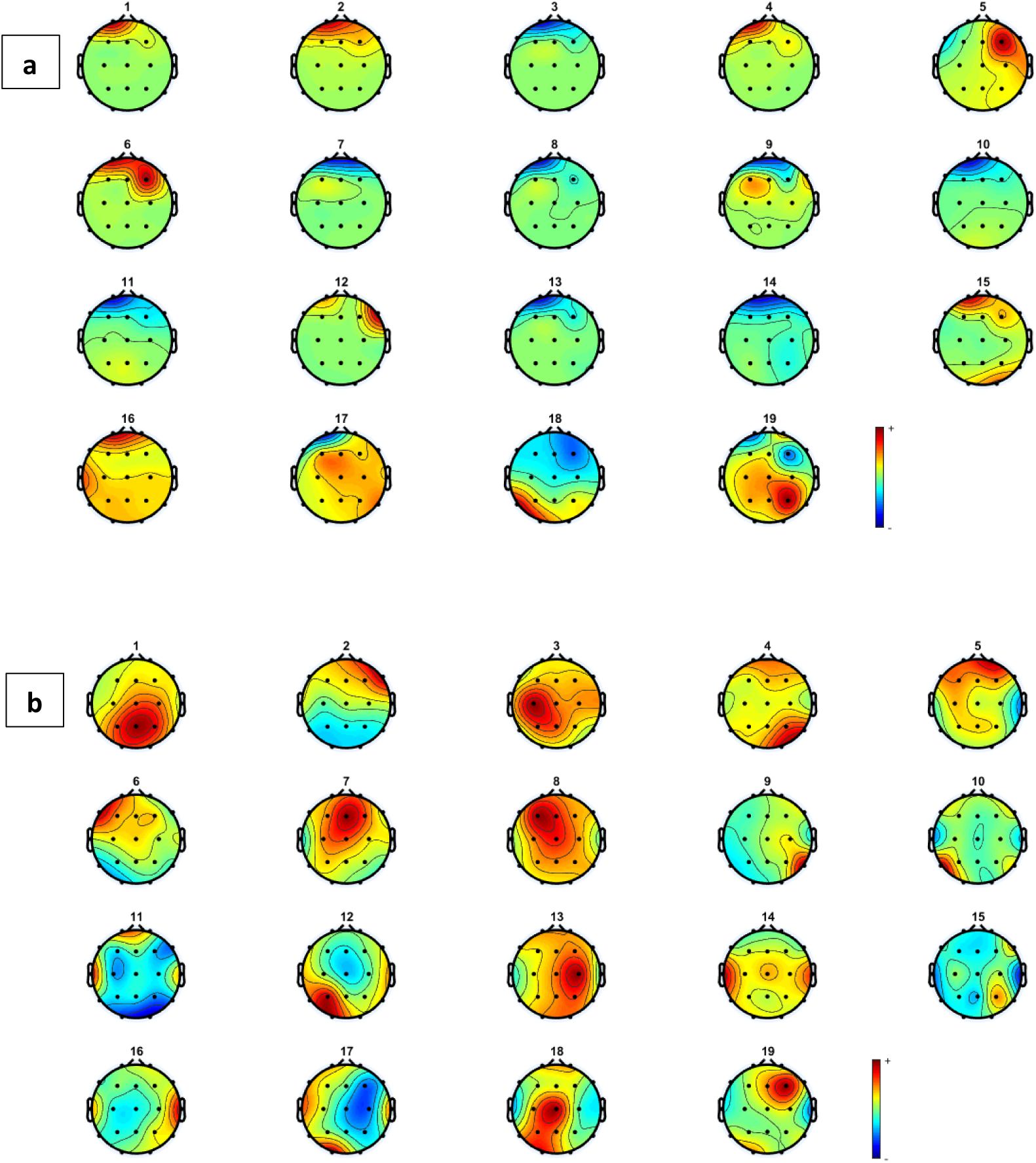
Topography maps for components related to (a) contaminated and (b) reconstructed and cleaned EEG.

**Fig 19.**
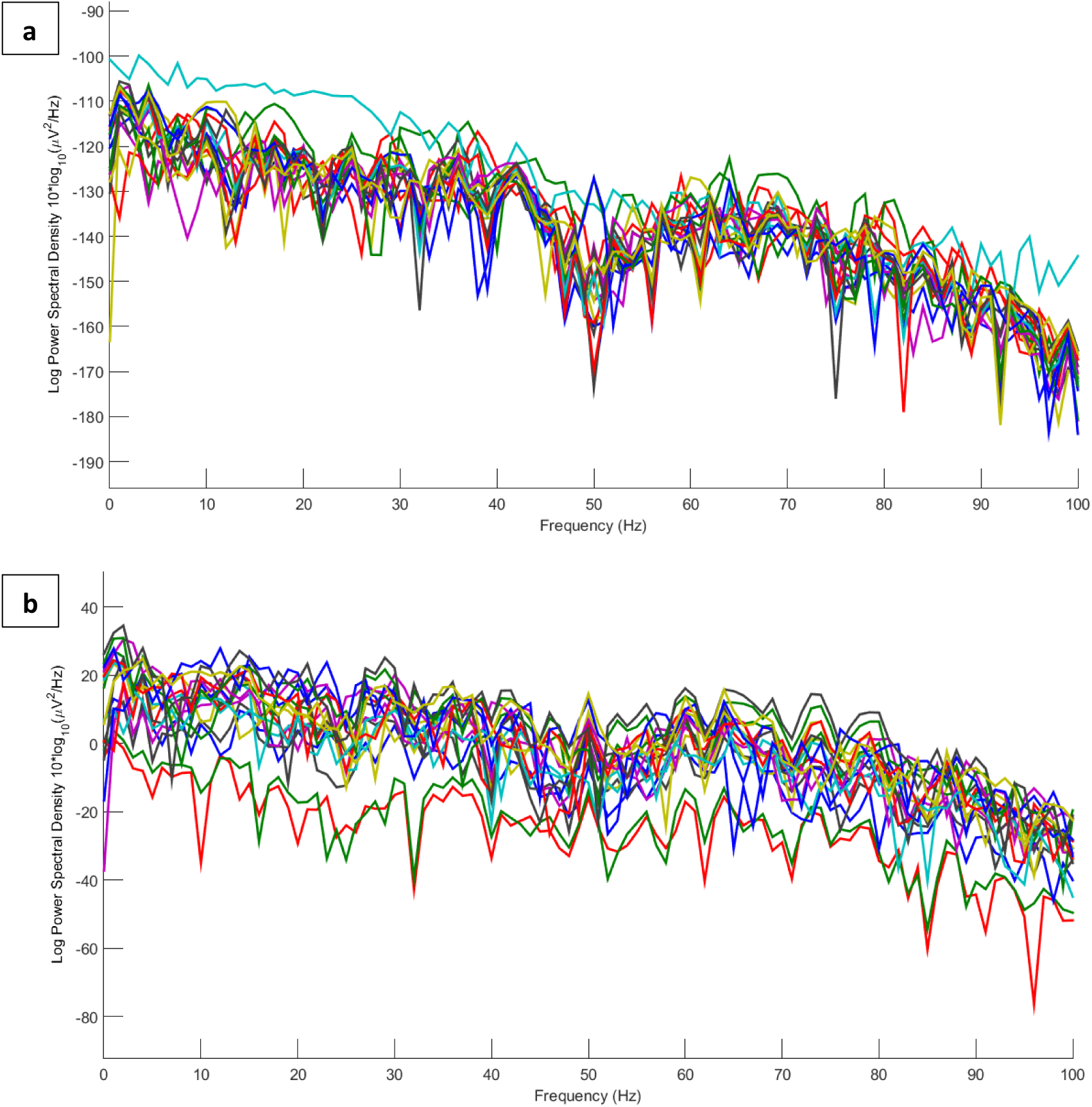
Power spectral density of channels for (a) contaminated real and (b) cleaned EEGs

Considering the results in Fig 19, artifacts are easy to distinguish in most components. Eye movement and blinking are clear with respect to the channel locations. The 12th component, for example, shows the activity in both sides of forehead which is related to ECG and can be seen in F8 channel. The topography map for components after artifact removal ensure the proposed method to work effectively. Hot spots are easy to see in topography maps before artifact removal. Fig 20 shows the power spectral density for all channels before and after the proposed method.

**Fig 20.**
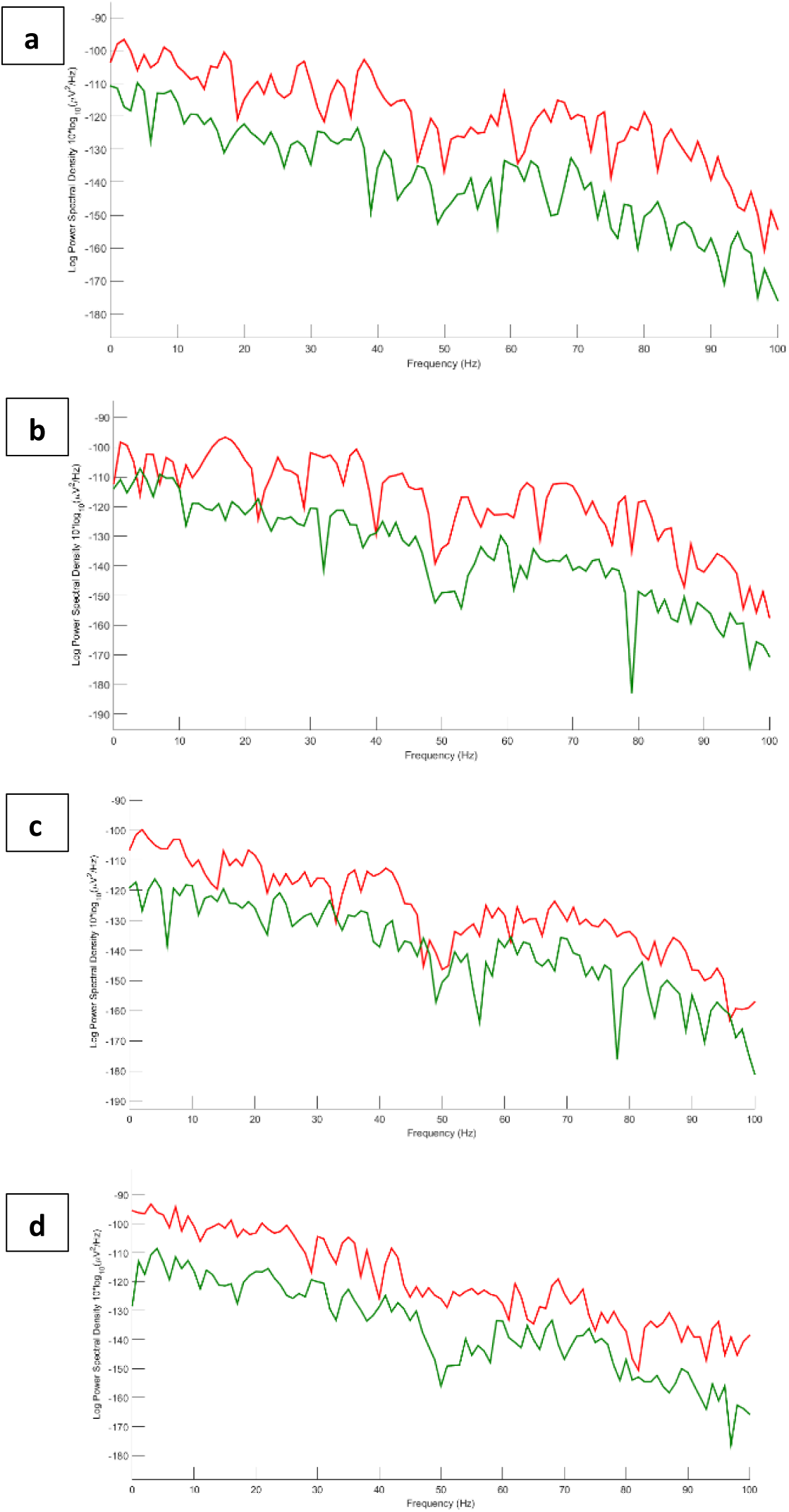
Power spectrum for (a) Fp1, (b) Fp2, (c) F8 and (d) T7. Red and green lines correspond to EEGs before and after applying artifact elimination respectively.

We also decide to study the results in frequency domain more comprehensively. Therefore, four channels contaminated by severe artifacts are selected and power spectral density for those channels are represented in Fig 21. These five selected channels contain Fp1, Fp2, F8 and T7. Red and green colors denote EEGs before and after applying the proposed method respectively.

**Fig 21.**
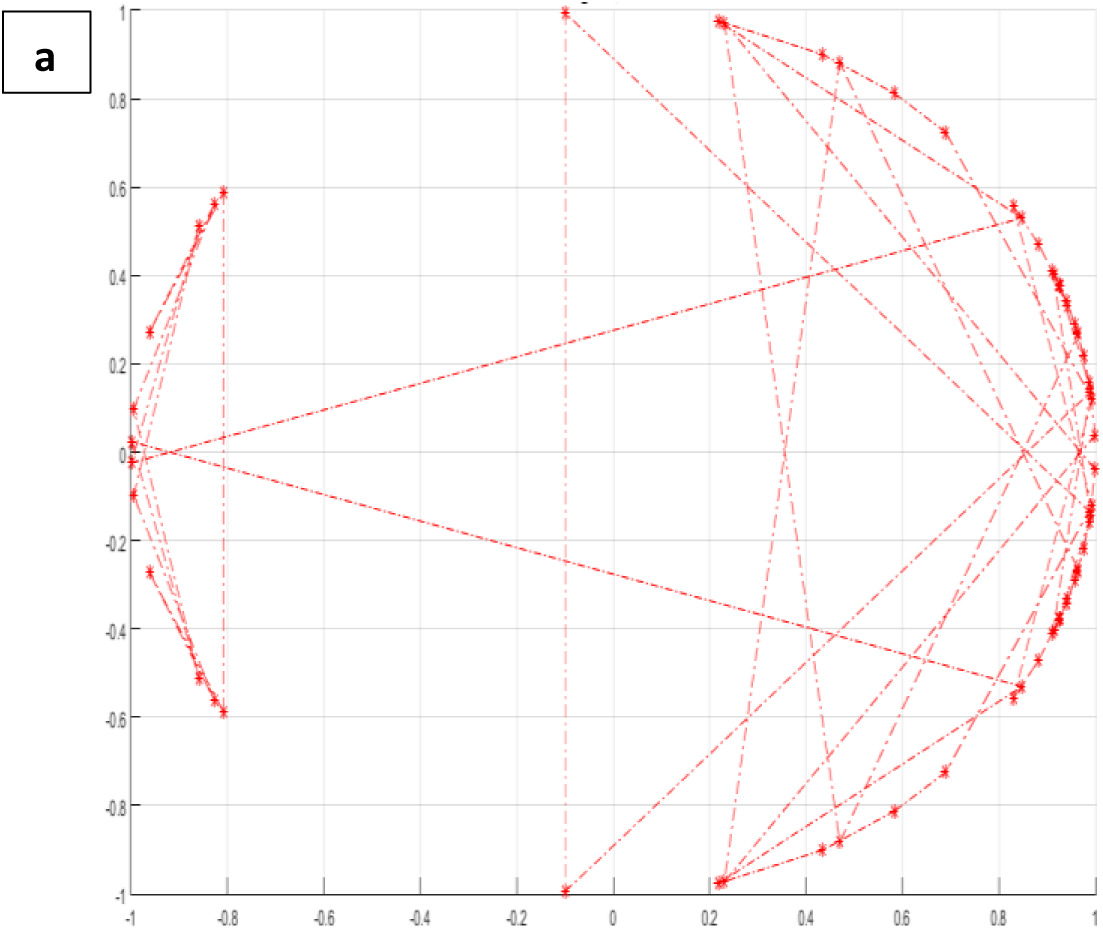
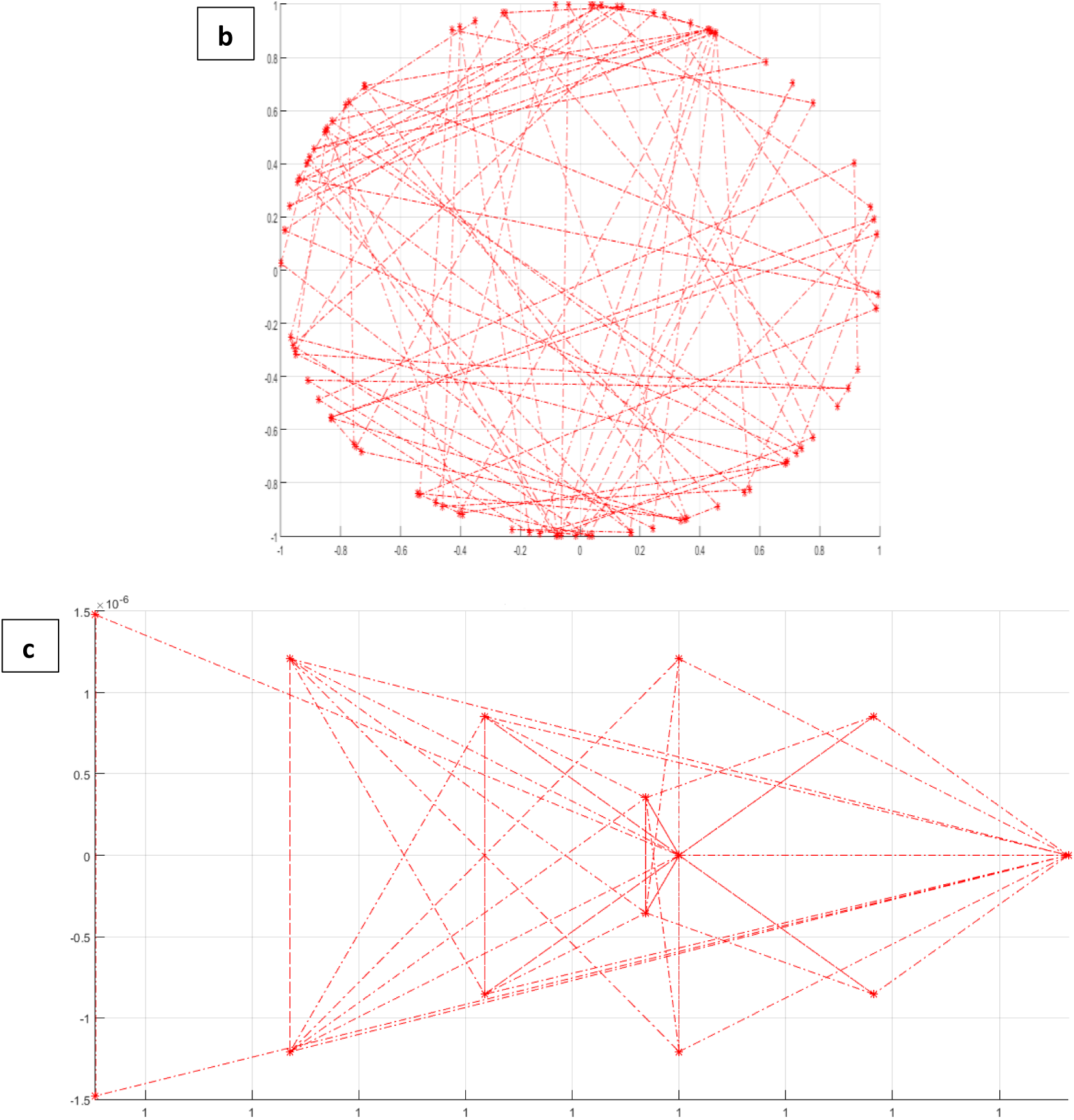
angle plot for (a) sinusoid (b) random (c) chaotic signals.

**Fig 22.**
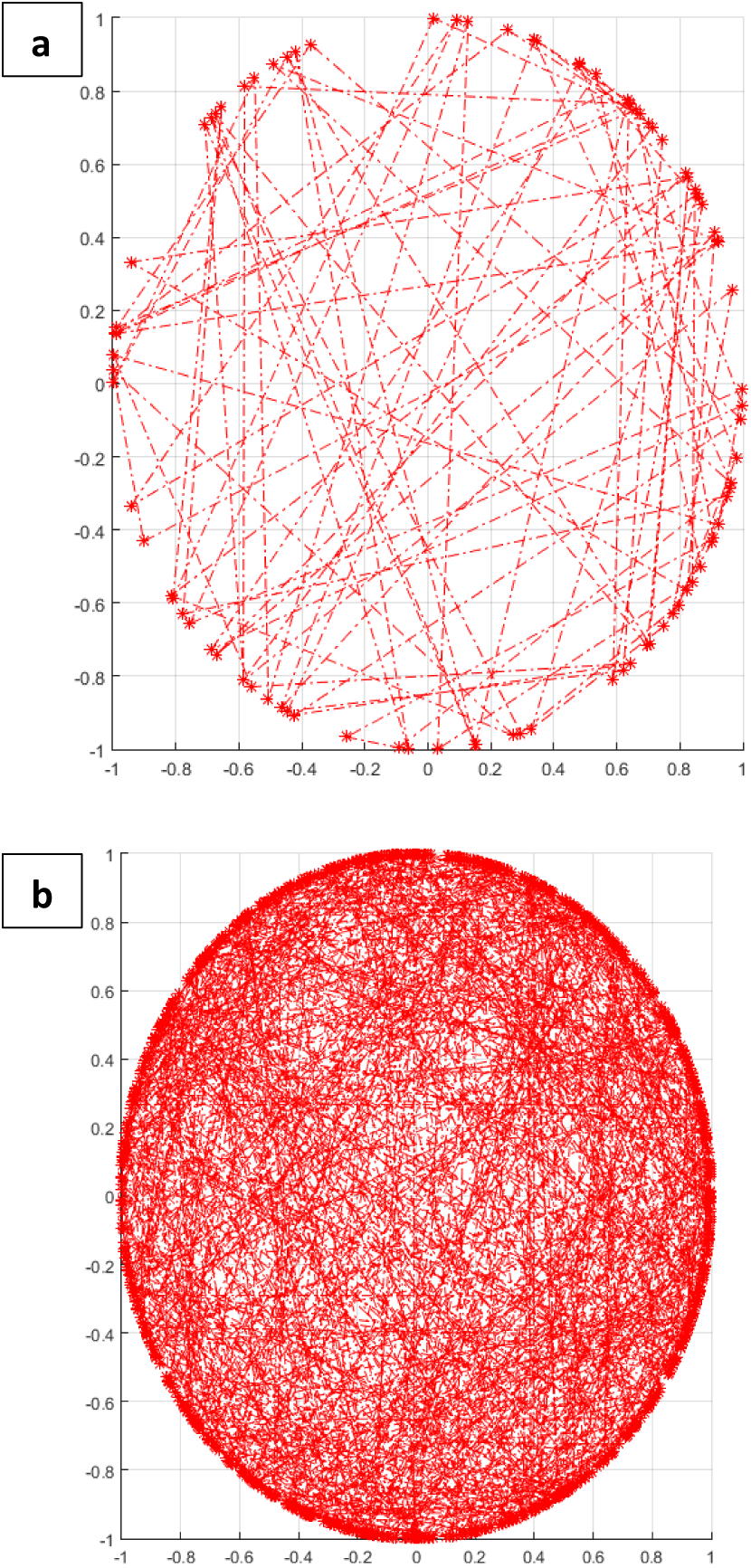
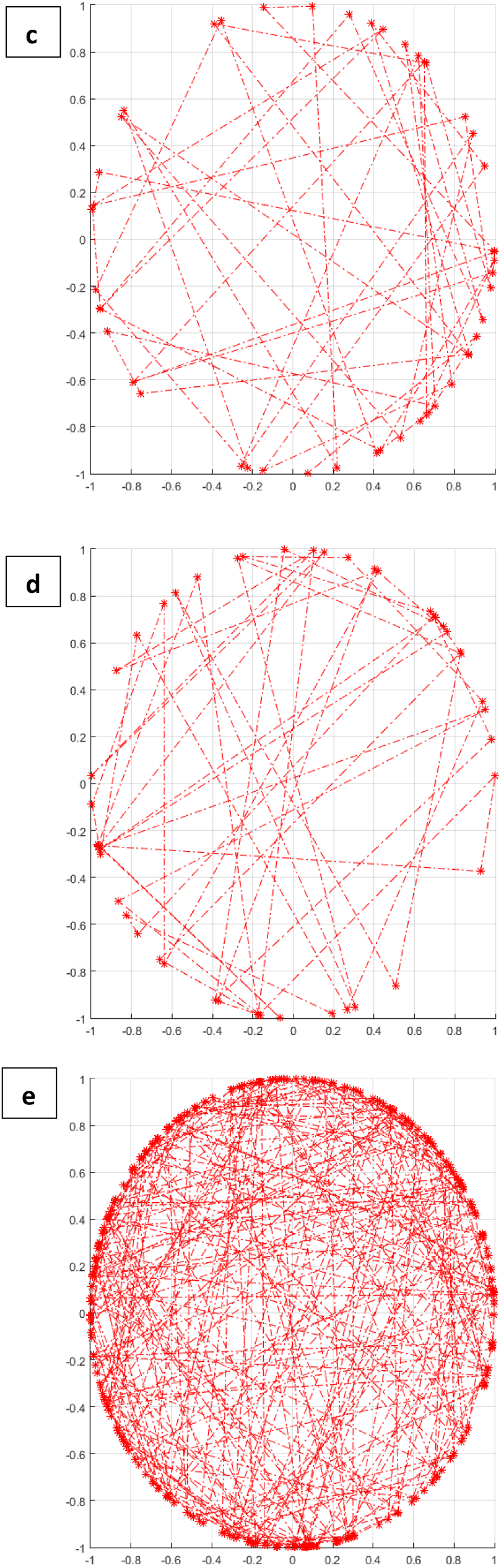
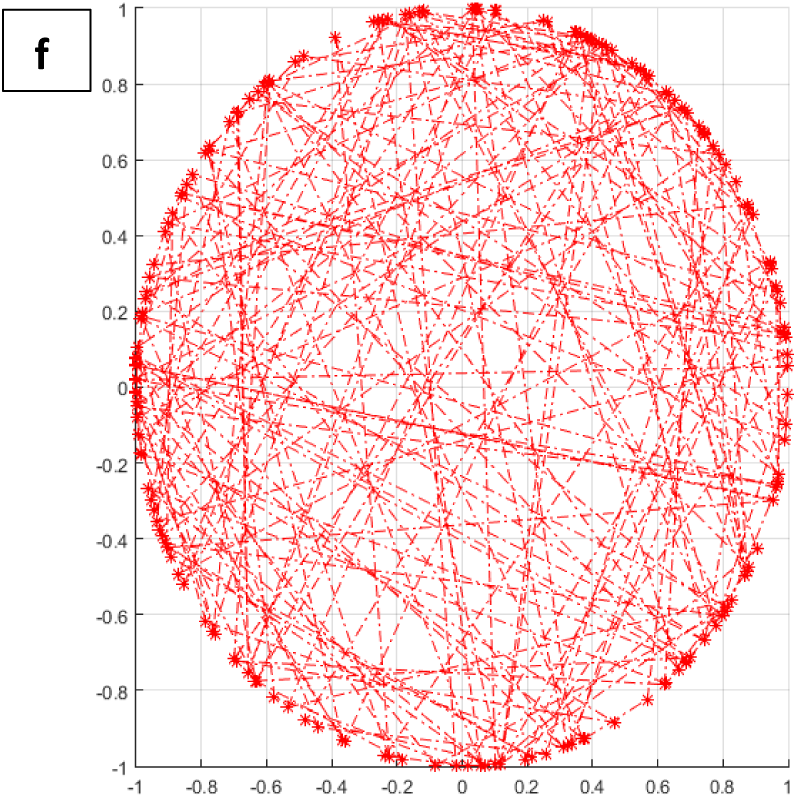
angle plots for single channel artifactual EEGs severely contaminated by (a) ECG, (b) EMG, (c) EOG, (d) eye blinking and (e) white noise. (f) angle plot for artifact-free EEG. All signals are sampled at 256 Hz and have the length equal to 10 s.

**Fig 23.**
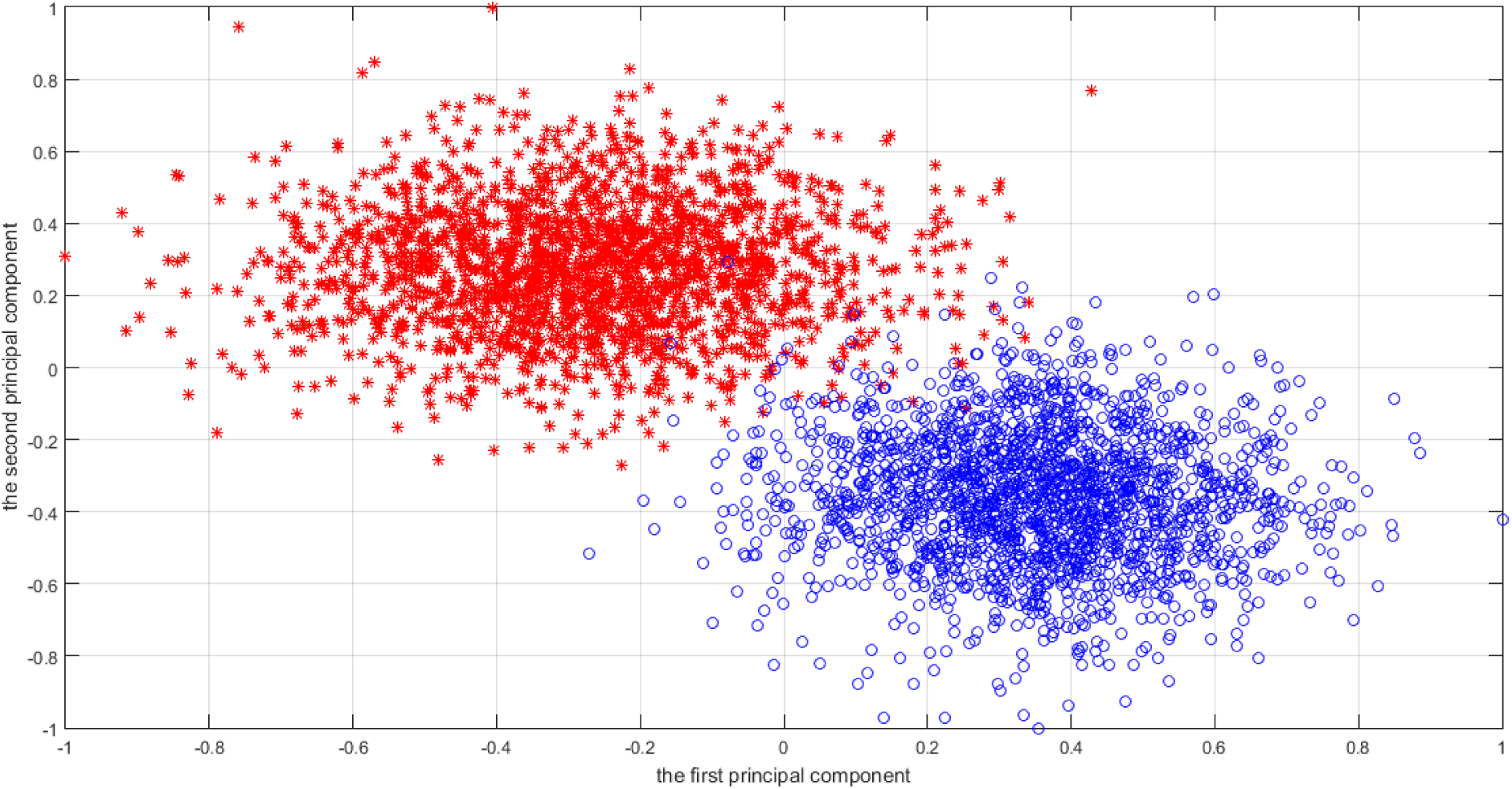
Scatter plot of the two largest principal components of the feature space over all datasets.

## 4. Discussion

In this study, we have managed to suppress different kinds of artifacts through an automated procedure. Conventional methods usually take advantage of filtering frequency bands which is not effective and practical since there is an overlap between EEG and most artifacts and noise in the frequency domain. Several surveys [2, 3, 4, 10, 11 and 12] have come to a conclusion that ICA-based methods can solve this problem. In [2 and 4] most BSS methods are evaluated at different levels of SNR and comparable results are achieved in those studies. On the other hand, these methods need artifactual sources to be determined by experts or classifiers which means that they are semi-automated. This motivates us to introduce a new automated artifact recognition system in the current study. Additionally, in some studies like [53], some features such as signal amplitude or slope are considered to recognize artifacts which is not appropriate and practical in most real cases. Due to the presence of noise and also different EEG recording systems and procedures, these measures require to be set for every single recording. In [50 and 54], authors have used correlation coefficients to sort extracted sources. In that study, the first and the last sources are considered artifactual and related to EOG and EMG which might result in information loss. Since we are not aware of the number of artifactual sources, in some cases there might be different numbers of artifactual sources from different types. These measures such as correlation coefficient and mutual information are not able to distinguish and recognize all components. Additionally, some sources are combinations of two or three artifacts which is why they cannot be completely recognized using just one or two criteria. In addition, there might be other types of artifacts such as those pertaining to eye movement or the white noise. So it seems that the proposed method in [54] is not very useful in most practical and real situations. To the best of our knowledge, there is no global measure or feature based on which artifactual sources can be recognized very accurately. That is the reason why we try to introduce an automated artifact recognition method based on the components’ dynamics in this study. Based on [50], BSS methods can effectively isolate brain activity and artifacts. Therefore, components are extracted based on one of best BSS methods which is called SOBI and is one the most common and effective source separation methods. Moreover, in numerous studies such as [2, 3, 4, 9, 10, 50] SOBI is claimed to be one of most effective and successful source separation methods which is able to isolate artifactual components. SOBI can effectively separate neural activities and artifacts [2 and 4]. Time-lagged covariance matrices are iteratively used in SOBI while separating sources. In addition, SOBI reduces computation complexity and is simple to implement [2, 3, 9 and 10]. Therefore SOBI is applied in this study. Because of EEGs and artifacts being non-stationary and nonlinear, we decide to examine the phase space of the components in order to recognize artifact ones and eliminate them. Several studies have proven of equivalency of phase space and the original system according to topological characteristics. In other words, phase space is likely to provide more and / or different information in comparison with common feature extraction methods [43]. It is believed that signal phase is of great importance while processing non-stationary and nonlinear signals and more information is embedded in signal phase in comparison with the signal magnitude. This is our motivation to go for the phase space of the components and propose a new state space called angle space. Angle space is capable of representing components’ dynamics. To study the properties of the angle space more completely, we decide to reconstruct the angle space for three different signals containing a sinusoid, a random and chaotic time series. Fig 20 represents the angle plot for these signals. It is evident that angle plots are significantly different. These signals are sampled at the same sampling rate and have the same length.

The sinusoid signal has the frequency equal to 10 Hz. The random time series is zero-mean with unit variance. The chaotic signal is achieved employing the logistic regression with the tuning parameter equal to 3.9. To investigate more, we decide to reconstruct the angle plot for all types of artifactual components. These angle plots are shown in Fig 21. Expectedly, angle plots are significantly different and can be recognized visually. For further analysis, we should describe the angle plot using some measures.

As it is clear, we can easily distinguish different types of artifacts visually in Fig 21. Some features and quantifiers are required to present the angle space mathematically. Poincare planes are applied to quantify the angle space of components. Besides, some statistical features are extracted from angle space. It is shown that extracted features from a higher dimensional transformations are more significant and discriminant than those in EEGs. Based on the results in table 5, most extracted features are significant and effective to distinct neural activity and artifacts. Three common classifiers, including MLP, KNN and Bayes, are employed to classify components. We also employ the mixture of the mentioned classifiers which is more accurate based on the achieved results. Experts have labeled all extracted sources and also monitored all recordings and simulations. Results show that the mixture of those three classifiers is more accurate while recognizing neural and artifactual components. As mentioned before, accuracy while classifying components into neural activity and five different types of artifacts is high enough and comparable to previous studies. Artifact sources, recognized by the classification model, are fed into SWT-based artifact elimination procedure which is introduced and evaluated in [54]. SWT has some remarkable characteristics in comparison with DWT and CWT which are explained before. In previous studies, all components are thresholded using wavelet transform or artifact ones are set to zero. Both approaches might pose serious problems especially in real BCI applications with large number of channels. This fact is also mentioned in several studies [10, 53, 54]. In this study, we are able to detect artifactual sources and eliminate artifacts using SWT. This suggested approach leads to better results in comparison with conventional methods which mostly result in desired information loss. In [54], several denosing methods are compared and based on the results, it is mentioned that eliminating artifacts using SWT is more efficient than threshloding all components using wavelet or setting artifact sources to zero. That is why we go for SWT-based artifact removal. In contrast to several studies such as [50, 53, 54], the proposed method in this study is completely automated and more precise. Since previous studies use different performance criterion and measures, it is impossible to compare their results with that of the current study. But we have tried to compute major evaluation parameters in order to have a fair comparison. In addition, previous studies have applied their methods to different datasets. So it seems to be difficult to compare the results. In terms of computation complexity, it should be noted that it takes the proposed method less than 0.25 s to analyze a 1-min and 19-channel EEG recording sampled at 256 Hz and remove artifacts. For all simulations and recordings, the processing time is under 0.25 s which is practical for BCI applications and also diagnosis purposes. All implementations are performed using MATLAB (release R2016a) running on Windows 7 Laptop PC with Intel(R)Core(TM)2 Duo 2.0 GHz processor with 4 GB RAM. The average processing time and the standard deviation for real EEG recordings and simulated EEGs are 0.21 s and 0.09 s respectively. Since other similar methods are compared with SWT-based artifact removal in [54], we avoid to review them here for sake of space. In this study, since only artifactual components are fed into SWT-based artifact removal procedure, it is clear that the proposed method has less computation complexity in comparison with methods which analyze all components.

Table 10 shows the average processing time and the standard deviation for all components at signal length of 10 s, 30 s and 60 s. There might be other BSS methods suggested in some studies like [50] which claims that some other BSS methods outperform SOBI in particular situations but while considering all evaluation measures such as processing time and simplicity, it is obvious that SOBI is slightly better than most BSS methods.

**Table 10.**
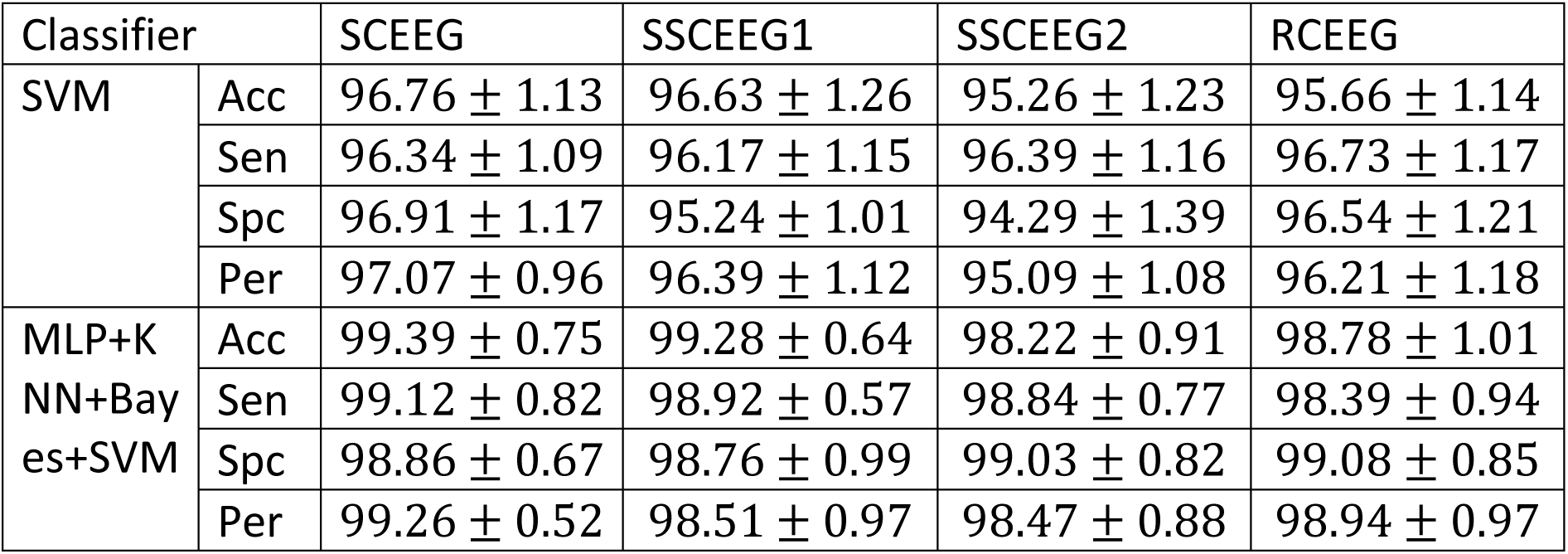
Classification accuracy of all the classifiers and also the mixture of them. Average values are mentioned here.

Since support vector machine (SVM) is claimed to be one the best and most common classifiers in the previous studies like [67], we decide to employ SVM with the polynomial kernel to classify components. Extracted sources are classified into two groups named brain-activity and artifacts. The SVM’s kernel and also other parameters are set based on trial and error. SVM Classification results are no better than other mentioned classifiers. SVM has almost the same results since the average accuracy is somewhat equal to other classifiers. The reason is mainly related to the separability in the feature space. Ten-fold cross validation is performed to evaluate SVM. Results show that the mixture classification model can slightly better recognize components. One can say that SVM can also be included in the mixture. This motivates us to take advantage of SVM in order to build the mixture classification model with the aim of artifact recognition. Table 12 represents the classification results. It is worth mentioning that components are classified into “brain-activity” and “artifacts” groups. Ten-fold cross validation is carried out.

For sake of space and also due to simplicity, we decide to bring average values for 10 s, 30 s and 1-min components in each dataset. It should be noted that we use ensemble voting to build the mixture of the classifiers. Results in table 12 suggest that the proposed mixture is able to recognize artifacts and separate them from brain activity. It is worth mentioning that the final mixture of experts which includes SVM, is more reliable and accurate than other proposed classification models. So we can realize that this mixture can be performed in future to classify artifacts and neural activity. Additionally, SVM’s results are slightly better than MLP and Bayes but statistical analysis shows no difference between classifiers while used alone. In contrast to single classifiers, the mixture classification models have better results and higher accuracies. Based on t-test, both mixtures (MLP+KNN+Bayes and MLP+KNN+Bayes+SVM) have significantly higher results than other classification models. As it is mentioned, we apply the voting between classifiers which means the test sample falls into the class with more votes. In terms of 3-classifier mixture, one class easily has more votes but in 4-classifier mixture, if the votes are equal, we go for SVM’s vote since it has slightly better results than other classifiers. It should be noted that voting is one of the simplest and fastest methods to fuse different classifiers.

To investigate more, we prefer to analyze the feature space more. To do that, all components from all datasets are normalized and then 12 suggested features are extracted from each source. We perform principal component analysis over all samples from different datasets and then normalize the components to achieve main and important components from the feature space. The two first components are plotted in Fig 21. Red and blue circles indicate artifactual and neural components respectively. 2000 samples are selected randomly from each class i.e. neural activity and artifacts to have equal number of samples in each class.

It is clear that feature space is separable. It can be verified that the proposed features can efficiently determine artifactual and neural components in this study. As we mentioned before, contaminated EEGs at different signal length are analyzed through the proposed method. Results show that although classification performance is the same for almost all components at different length, 10-s components are better classified in comparison with components with the length of 30 s and 60 s. As it is mentioned before, the phase space and consequently the angle space are able to demonstrate and represent the signal dynamics even at short length of the signal. This nonlinear analysis provides us with some new features which do not vary based on the length of the signal. That is why we can recognize sources very well regardless to the signal length. We have also tried different sampling rates. Real and simulated signals are sampled at the sampling rate of 128, 256, 512 and 1024 Hz. No significance difference is found in the results between different sampling rates. Similar results at different signal length and sampling rates suggest that the proposed method is quite effective and practical for different purposes such as BCI and diagnosis. Also it should be noted that all results are acceptable and comparable in terms of visual subjective inspection and also quantitative objective measures. It is worth mentioning that that in this study we focus on removing stereotyped biological artifacts. Non-stereotyped artifacts such as head and electrode movement might cause special pattern while recording EEG. These artifacts should be eliminated before the proposed artifact removal procedure. Fortunately, these artifacts can be easily discarded from the data by visual inspection.

In term of computer simulations, it should be mentioned that all simulations regarding the proposed method have been implemented in MATLAB. We also take advantage of EEGLAB [REF] in order to separate and analyze sources.

## 5. Conclusion

In this paper, we introduce a new method to suppress different types of artifacts and noise based on BSS (SOBI), wavelet transform (SWT) and a mixture of classifiers (MLP, KNN, Bayes and SVM). A preprocessing chain is suggested and evaluated in this paper. Results suggest that the proposed method can be employed to eliminate EEG artifacts. We have come to a conclusion that the proposed method is effective, fast and simple. Based on the results in this study and previous suggestions, hybrid methods which include BSS methods and artifact elimination procedures are recommended in order to correctly remove artifacts and noise from EEG [10]. Single channel artifact removal methods such as wavelet transform have been widely employed in most recent studies [2, 3, 10, 11]. It is mentioned and proved in [3] that wavelet transform should be a posterior step to BSS methods. It is also mentioned that automated methods are superior to methods based on visual inspection in terms of artifact elimination and EEG interpretation.

SOBI is based on independent component analysis which means that we should take some assumptions into consideration. In other words, some assumptions have been fulfilled in this study. These assumption are as following:

- component projections are mixed and summed linearly at channels,
- components are independent and
- components have non-gaussian distribution.

Considering artifacts and EEG, it is no exaggeration to say that the mentioned assumptions are realistic in most recordings and simulations in this study. Several studies like [53], have considered these assumptions and proved that linear decomposition methods such as SOBI seem to be appropriate to separate EEG components. Similar to most previous studies in this field, these assumptions are considered in this study. On the other hand, there are several advantages to employing BSS methods like SOBI in EEG analysis. SOBI allows direct examination of components in term of dynamics relate to separate or related regions in brain. In addition, most stereotyped artifacts are isolated by SOBI in different components.

Another important point is about low-dimension EEG signals. In this study, it is assumed that the number of sources is equal to or less than the number of channels. Therefore, sufficient number of EEG channels is required to correctly estimate sources. Moreover, it is considered that the number of artifacts is less than or equal to the number of components and channels. These assumptions might cause problems while dealing with low-dimension EEG signals. In that situation, some decomposition methods such as empirical mode decomposition might be a good solution to decompose EEGs as the first step. Then BSS methods can be applied to decomposed signals.

Since the mixture of several classifiers is employed in this study, it should be noted that over-fitting and under-fitting might cause some problems. It can be considered one of the weak points while applying classifiers in automated artifact recognition methods. This problem could be tackled by considering training and testing errors together and also performing some validation methods such as k-fold cross validation.

It is worth mentioning that, no global measure is available to evaluate and compare different methods in this field. Besides, previous studies have tested methods on different datasets and simulations. This make the results inconsistent. That is why most previous studies have trouble reproducing other methods. In this study, we have tried to evaluate the proposed method through several approaches. Temporal, spectral performance parameters are considered in this study. We will try to define new evaluation criteria in future work. It should be noted that, the results reveal the fact that although the proposed method outperforms most previous studies and is quite fast, effective and practical, it fails in a few cases while dealing with highly-contaminated EEGs. This motivates us to make attempts to introduce new methods in future. The proposed method has been applied to real, semi-simulated and simulated EEGs in order to remove artifacts in BCI applications. In future, this method could be applied to ictal and inter-ictal EEGs for diagnosis purposes and EEG interpretation. In future work, we are going to employ and evaluate the proposed method in real time BCI applications. Other BSS methods, classifiers and artifact elimination procedures could also be introduced and compared with this study. In addition, the proposed method could be employed to eliminate other types of artifacts such as power-line interference and head movement. However, these purposes are beyond the scope of the current study.

## Acknowledgements

The authors are thankful to …

